# Coding and noncoding somatic mutations in basal cell carcinoma

**DOI:** 10.1101/807313

**Authors:** Maria Giovanna Maturo, Sivaramakrishna Rachakonda, Barbara Heidenreich, Cristina Pellegrini, Nalini Srinivas, Celia Requena, Carlos Serra-Guillen, Beatriz Llombart, Onofre Sanmartin, Carlos Guillen, Lucia Di Nardo, Ketty Peris, Maria Concetta Fargnoli, Eduardo Nagore, Rajiv Kumar

## Abstract

Basal cell carcinoma (BCC) represents the most commonly diagnosed human cancer among persons of European ancestry with etiology mainly attributed to sun-exposure. In this study we investigated mutations in coding and flanking regions of the *PTCH1* and *TP53* genes and noncoding alterations in the *TERT* and *DPH3* promoters in 191 BCC tumors. In addition, we measured CpG methylation within the *TERT* hypermethylated oncological region (THOR) and transcriptions levels of the reverse transcriptase subunit. We observed mutations in *PTCH1* in 59% and *TP53* in 31% of the tumors. Noncoding mutations in *TERT* and *DPH3* promoters were detected in 59% and 38% of the tumors, respectively. We observed a statistically significant co-occurrence of mutations at the four investigated loci. While *PTCH1* mutations tended to associate with decreased patient age at diagnosis; *TP53* mutations were associated with light skin color and increased number of nevi; *TERT* and *DPH3* promoter with history of cutaneous neoplasms in BCC patients. *TERT* promoter mutations but not THOR methylation associated with an increased expression of the reverse transcriptase subunit. Our study signifies, in addition to the protein altering mutations in the *PTCH1* and *TP53* genes, the importance of noncoding mutations in BCC, particularly functional alterations in the *TERT* promoter.

## Introduction

Basal cell carcinoma (BCC) accounts for about 80 percent of all skin cancers and it is the most commonly diagnosed neoplasm among the Caucasian population^1-3^. The tumor originates from stem cells within hair follicles or the interfollicular epidermis and infundibulum^4,5^. Basal cell carcinoma rarely metastasizes; however, due to sheer number of people affected, the disease poses a considerable health hazard as it causes extensive morbidity through local invasion and tissue destruction^6^. The most common genetic alterations in BCC involve the Hedgehog signaling pathway. Germline mutations in the Hedgehog receptor patched 1 (*PTCH1*) occur in patients with Gorlin syndrome, a Mendelian disease with a high prevalence of BCC^7^.

Aberrant activation of the Hedgehog pathway is common in sporadic BCC either through mutations in the *PTCH1* gene, activating mutations in smoothened (SMO) or loss-of-function mutations in suppressor of fused homolog (SUFU)^8-10^. Ubiquitous hyper-activation of the Hedgehog has led to development of inhibitors targeting G-protein coupled receptor Smoothened for treatment of advanced BCC^9,10^. While, the studies based on exome sequencing have confirmed the centrality of *PTCH1* mutations in BCC, the variants within smoothened are reportedly causal in resistance to the inhibitors^11^. The role of other frequent genetic variants in either tumor development or resistance to treatment cannot be ruled out. Other prevalent alterations in BCC include loss of function mutations in *TP53*^12^.

In addition, various reports have suggested frequent occurrence of mutations within the promoter region of the gene encoding telomerase reverse transcriptase (*TERT*), the catalytic subunit of telomerase^13^. The mutations within the *TERT* promoter create *de novo* binding sites for ETS transcription factors that lead to increased transcription through massive epigenetic histone modification^14,15^. Methylation within *TERT* hypermethylated oncological region (THOR) also reportedly de-represses the *TERT* transcription^16^. Besides *TERT* promoter, other frequent noncoding mutations in BCC include those involving a bidirectional *DPH3* promoter^17^.

In this study, we sequenced 191 BCC lesions and corresponding apparently normal skin surrounding tumors from 115 patients for mutations in the *PTCH1* and *TP53* genes, and the *TERT* and *DPH3* promoters. We also investigated CpG methylation at the *TERT* promoter between -560 to -774 bp region from the ATG start site that falls in the THOR. We also measured the effect of the *TERT* promoter mutations and THOR methylation on the transcription of the catalytic subunit of the telomerase. In addition, we also investigated the correlation between *TERT* promoter mutations and telomere length.

## RESULTS

### PTCH1 mutations

We detected 137 *PTCH1* mutations in 105 tumors; of those 44 tumors also showed loss of heterozygosity as detected by MLPA including focal deletions on one allele in 4 tumors (**Table 1; Supplementary Figure 1; Supplementary Table 1 and 2**). In addition 7 tumors showed only loss of heterozygosity without a mutation on the remaining allele. One mutation each was detected in 79 (41%) tumors, 21 (11%) tumors carried two mutations each, four tumors had three mutations each and one tumor had four mutations.

**Table 1:**
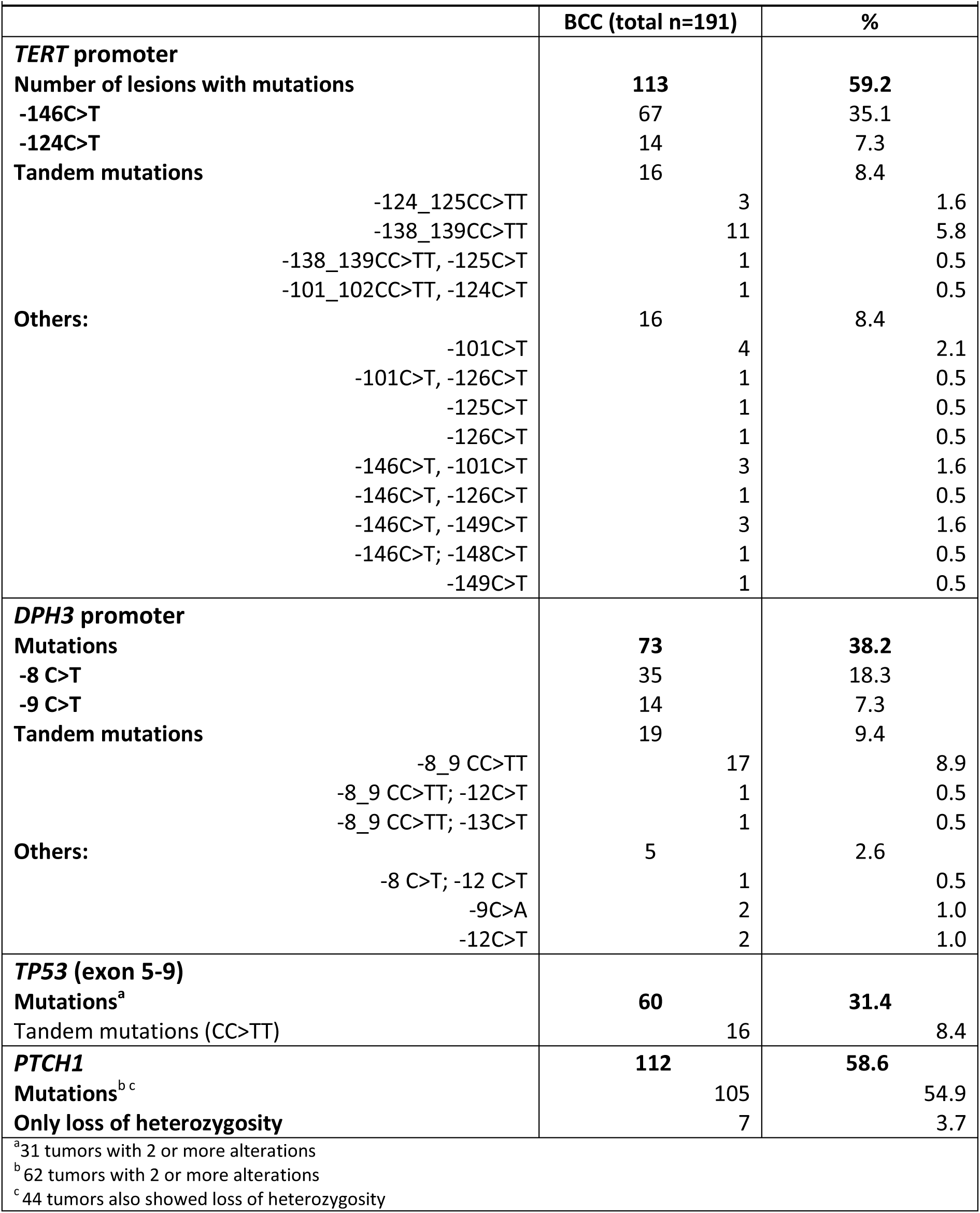
Mutations in *TERT* promoter, *DPH3* promoter, *TP53* and *PTCH1*.

The C>T base change was the most frequent mutation with 64 transitions detected in 56 (29%) tumors, followed by CC>TT tandem mutations, and C>A mutations in 8 tumors each. Thirty-six mutations were missense, 33 were nonsense and 17 mutations were intronic (**Figure 1)**. We also detected 37 insertion-deletions (indels) that included 25 frame-shift alterations, 6 in-frame deletions, two truncations, and three intronic. Out of three duplications, two were frame-shift and one in-frame. We also detected 14 mutations at 5’ splice sites and 7 mutations at 3’ splice sites (**Supplementary Table 3**). Mutations were distributed throughout the gene from exon 3 to exon 23; exon 23 with 12 mutations had the highest number of alterations followed by exons 18 and 6 with 10 mutations each. Most of the missense mutations were in exon 23 while exon 18 had the largest number of nonsense mutations and indels were most frequent in exon 20.

**Figure 1.**
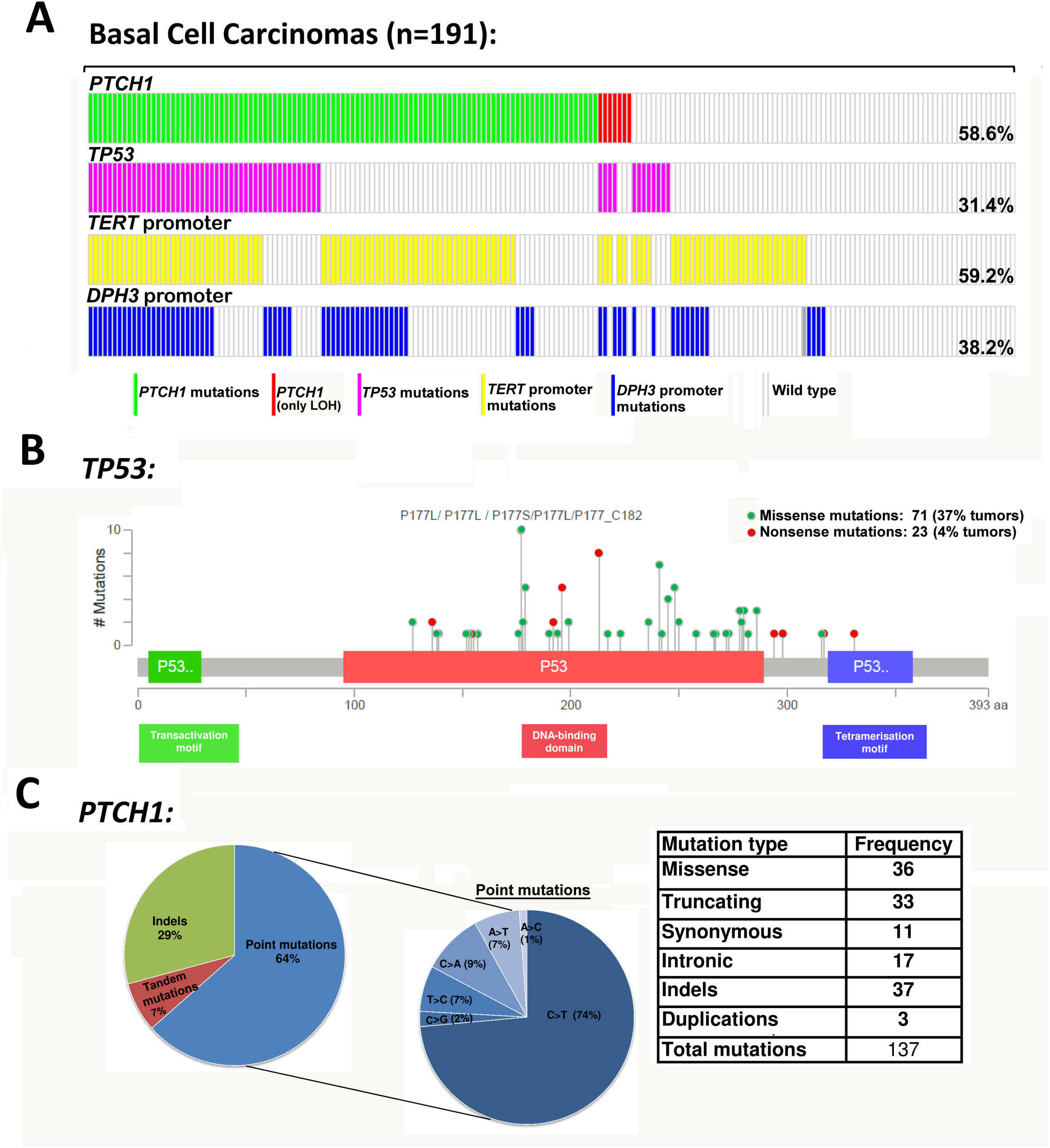
Mutations in skin basal cell carcinoma tumors: **A. Distribution of mutation at** *TERT* promoter, *DPH3* promoter, *TP53* and *PTCH1* gene. **B. Distribution of mutations within different p53 domains**. Protein diagram was generated with cBioPortal tools. **C. Frequency and type of mutations in *PTCH1* gene.**

### TP53 mutations

Sixty of the 191 (31%) BCC carried alterations in *TP53*, with 31 (16%) tumors carrying more than one mutation; 24 tumors carried two mutations and 4 tumors carried three mutations and 3 tumors carried four mutations. In total, 100 alterations in the *TP53* gene were identified. The most frequent mutations were C>T transitions with 66 transitions in 48 (25%) tumors, followed by 17 CC>TT tandem transitions in 16 (8%) tumors (**Table 1**; **Supplementary Table 1; Supplementary Table 4**).

The majority of *TP53* mutations were missense, 71 mutations in 51 tumors (37%) and 23 nonsense mutations were present in 20 tumors (4%). The mutations were distributed within the DNA binding domain of *TP53* (**Figure 1**). The most common missense variant p.P177L (c.530 C>T) in exon 5 was detected in 8 tumors (4%) followed by the p.H179Y (c.535 C>T) mutation in exon 5 in 4 tumors. The truncating mutations detected with the highest frequency were the p.R213* in exon 6 (c.637 C>T) identified in 8 (4%) tumors and p.R196* in exon 6 (c.586 C>T) in 4 (2%) tumors.

### TERT promoter mutations

*TERT* promoter mutations were present in 113 of 191 (59.2%) lesions with the -146C>T mutation in 67 (35.1%) tumors followed by -124C>T in 14 (7.3%) tumors (**Table 1; Supplementary Table 1**). CC>TT tandem mutations at -138_139, -124_125 and -101_102 bp positions were present in 16 (8.4%) tumors. Eight BCC tumors in addition to the -146C>T hotspot mutation also carried additional alterations and in five tumors other than the recurrent hotspot mutations were detected (**Table 1**).

### TERT promoter methylation

Pairwise alignment of bisulfite and target genomic sequence using QUMA showed > 85% of homology in 157 (86%) of 183 tumors. Of those 157 tumors, 93 (59%) carried *TERT* promoter mutations while the remaining 64 (41%) were wild type. The methylation at the screened 14 CpG sites was statistically significantly higher in tumors without the *TERT* promoter mutations (99%; 870 of 882 methylation sites) than in the tumors with those mutations (96%; 1219 of 1267 methylation sites; Fisher’s exact P: 0.0006, Mann-Whitney U-test P: 0.003). The standard errors (SE 0.65 vs 0.78%) between the two groups were not statistically significantly different (F test P: 0.12). Comparison of methylation at individual CpG sites showed a statistically significant differences at positions chr5:1,295,731, hg 19 (97% in tumors without mutations vs 80% in tumors with mutations; Fisher’s exact P: 0.003) and chr5:1,295,759 (98% in tumors without mutations vs 90% in tumors with mutations; Fisher’s exact P: 0.05).

Analysis of bisulfite converted DNA after cloning also showed statistically significantly higher methylation in tumors without mutations than in those with mutations (Fisher’s exact P: <0.00001, Mann-Whitney U-test P: <0.00001; **Supplementary Figure 2**). Pairwise sequence alignment of cloned bisulfite sequence and the target genomic sequence using QUMA showed more than 95% homology for each clone. Individually at the each CpG sites, tumors without the *TERT* promoter mutations had statistically significantly higher methylation than the tumors with mutations. The methylation across all 14 CpG sites ranged between 85% - 98% in tumors without mutations compared to 44% - 91% in tumors with mutations.

### DPH3 promoter mutations

Mutations in the *DPH3* promoter were present in 73 of 191 (38.2%) tumors. In addition to the frequent C>T transitions observed in 35 (18.3%) tumors at -8 bp and in 14 (7.3%) tumors at -9 bp, CC>TT tandem mutations at -8_9 bp were present in 17 tumors (8.9%). Additional mutations included two C > T transitions at -12 bp, and two C > A transversions at -9 bp. The C>T mutation at -12 bp in two BCCs co-occurred with -8C > T and -8/-9CC > TT mutations, respectively, and the C>T mutation at -13 bp in one BCC co-occurred with -8/-9 CC>TT (**Table 1; Supplementary Table 1**).

### Association of mutations with patient and tumor characteristics

Overall, 28 (14.7%) tumors carried alterations at all four loci; 37 (19.4%) tumors carried alterations at three loci (*PTCH1* and *TERT* promoter along with alteration at either *TP53* or *DPH3* promoter); 32 tumors (16.8%) had alterations in 2 genes (*PTCH1* gene and *TP53* or *TERT* promoter or *DPH3* promoter); 14 (7.3%) tumors had only *PTCH1* mutations, while 42 (22.0%) tumors that were wild type for *PTCH1* and carried any of the other alterations **(Figure 1)**. *PTCH1* mutations tend to co-occur with: *TP53* mutations (OR 7.69; 95% CI 3.38-17.46; *P*<0.0001), *TERT* promoter mutations (OR 3.84; 95% CI 2.08-7.07; *P*<0.0001) and *DPH3* promoter alterations (OR 5.09; 95% CI 2.56-10.12; *P*<0.0001) (**Figure 1**).

The presence of *PTCH1* mutations tended to associate with decreased patient age at diagnosis (OR 0.58, 95%CI 0.32-1.04, P 0.07); *TP53* mutations associated statistically significantly with light skin color (OR 2.13, 95%CI 1.13-4.00; P 0.02) and >50 nevus count (OR 2.66, 95%CI 1.03-6.87, P 0.04). Non-coding mutations in *TERT* promoter (OR 2.02, 95%CI 1.03-3.97, P 0.04) and *DPH3* promoter mutations (OR 2.25, 95%CI 1.10-4.57, P 0.03) were associated with history cutaneous neoplasms (**Figure 2; Supplementary Table 5)**. In the multivariate analysis, the association between *TP53* mutations and fair skin (OR 5.31, 95%CI 2.19-12.88; P 0002); *DPH3* promoter mutations and history of cutaneous neoplasm (OR 2.47, 95%CI 1.06-5.72; P 0.04) remained statistically significant. In a separate multivariate analysis the simultaneous presence of mutations at all four loci in BCC tumors associated with light skin color (OR 4.84, 95%CI 1.09-21.45; P 0.04) and history of cutaneous neoplasms (OR 8.85, 95%CI 1.51-51.80; P 0.02; data not shown).

**Figure 2.**
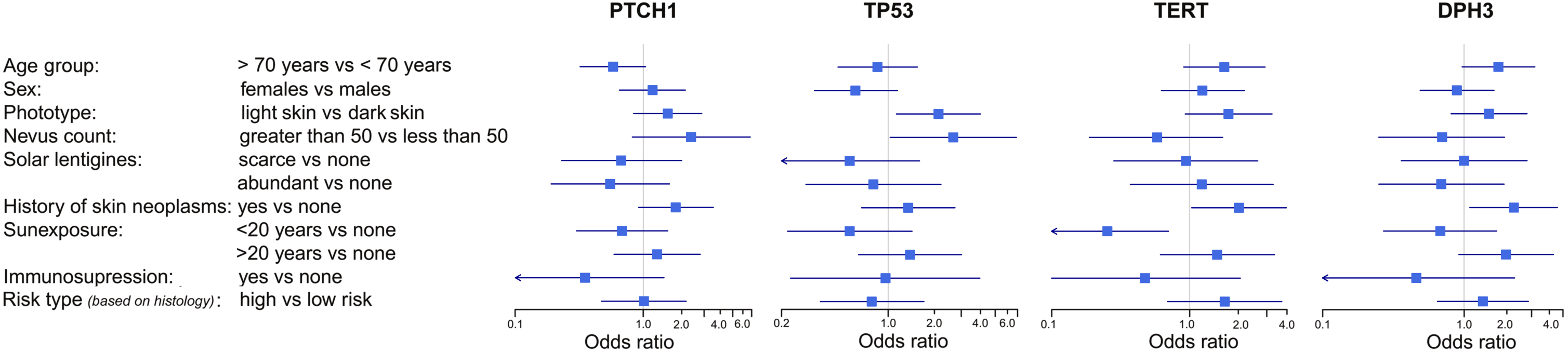
Association between mutations and tumor/patient characteristics: Forest plot were plotted to depict OR and 95%CI for the associations analyzed through univariate logistic regression.

#### *TERT* expression

RNA was available from 77 BCC tumors and corresponding tumor-surrounding skin. Of 77 tumors, 48 carried *TERT* promoter mutations, 34 with -146C>T, 8 with -124C>T, 6 with - 138_139CC>TT and 29 were without mutations. Data analysis showed statistically significantly higher mRNA levels in BCC tumors with *TERT* promoter mutations than in tumors without mutations (*P* < 0.001, *t*-test; **Figure 3**). Further stratification showed that tumors with *TERT* promoter mutations and complete methylation (n = 23) had statistically significantly (*P* = 0.003) higher *TERT* expression than the tumors with complete methylation and without *TERT* promoter mutations (n = 14) (**Supplementary Figure 3**). Similarly, the tumors with partial methylation and *TERT* promoter mutations (n = 25) had statistically significantly (*P* = 0.004) higher *TERT* expression than corresponding tumors with partial methylation and without the *TERT* promoter mutations (n = 2). The difference in *TERT* expression in tumors based on only methylation status was not statistically significant (*P* = 0.82).

**Figure 3.**
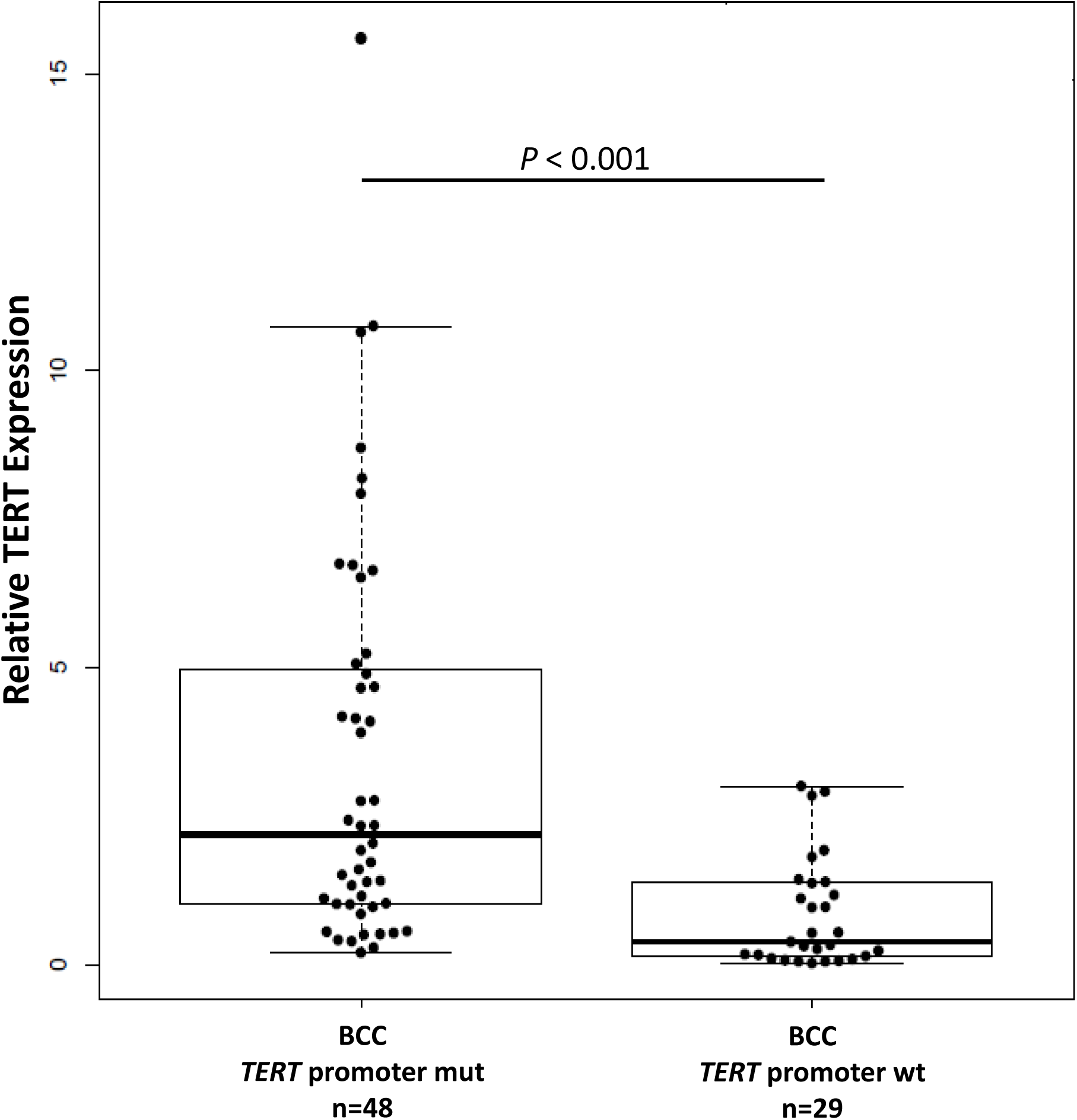
Relative *TERT* expression in BCC tumors based on the promoter mutational status: Experiments were carried out in triplicates and box plots represent mean ± standard error of means; *P*-value was determined by *t*-test.

#### Telomere length

Results from measurement of relative telomere length were available for 174 BCCs and 104 tumor-surrounding skin tissues. Of the 174 BCCs, 100 were with and 74 were without *TERT* promoter mutations. Relative telomere length ranged between 0.19 and 3.62 with median values of 1.02 in tumor-surrounding skin, 0.81 in *TERT* promoter wild type BCCs and 0.72 in BCCs with *TERT* promoter mutations. Tumors had shorter telomeres than surrounding skin; however, a difference in telomere length in BCCs with and without the *TERT* promoter was not statistically significant (**Supplementary Figure 4**).

## Discussion

We confirmed the presence of *PTCH1* and *TP53* mutations in BCC tumors, and showed frequent noncoding mutations within the *TERT* and *DPH3* promoters. *TERT* promoter mutations, the most frequent alterations, were associated with increased expression of telomerase reverse transcriptase subunit. The increase in *TERT* expression was due to the mutations and not due to hypermethylation of the oncological region within the *TERT* promoter.

*PTCH1* is a human homolog of the Drosophila segment polarity genes and its inactivation confers Sonic Hedgehog independent growth, genomic instability and tumor development potential. Hedgehog ligands bind to *PTCH1*, causing internalization and degradation leading to release of smoothened that in turn cascades the transcription of Hedgehog pathway genes in a feedback loop mechanism^18^. Inactivating alterations in *PTCH1*, consisting of 23 exons, were present in about 59 percent of tumors. In accordance with a central role of UV radiation in the pathogenesis of BCC, the majority of point mutations were represented by C>T and CC>TT transitions as also reported in previous studies^8,19,20^. The observed frequency of the *PTCH1* mutations in this study was similar to reported in earlier studies albeit lower than that reported in an exome sequencing based genomic analysis^8,12^.

The *TP53* genes encodes p53 transcription factor, a tumor suppressor involved in cellular stress responses including DNA damage, oxidative stress, oncogenic hyperproliferation, pathogenic stimuli, UV-induced pigmentation^21,22^. Approximately 31% of tumors carried mutations in the gene and a majority of mutations reflected UV signatures. Most of the base changes were predominantly protein altering missense changes and to lesser extent nonsense mutations. The mutations mainly affected the DNA binding domain with typical hotspot mutations as reported earlier in a study based on exome sequencing^12^. The selection for missense mutations in DNA binding domain the gene is driven through dominant negative effect where mutant forms hinder functioning of the protein from intact alleles^23^. The observed frequency of the mutations in our study was closer to that reported in non-aggressive than in aggressive BCCs^24,25^. However, in a study based on exome sequencing on a large series of BCC tumors, the frequency of mutations in *TP53* was reported to be 61%^8^. We also observed a strong association between *TP53* mutations and fair skin; impaired p53 in mice has been shown to result in lack of tanning response and addiction to sunlight^21,26^.

While the role of mutations in the tumor suppressors *PTCH1* and *TP53* is well established in BCC pathogenesis, the relevance of noncoding mutations is not well defined. Originally discovered in melanoma, the *TERT* promoter mutations lead to creation of *de novo* sites for ETS transcription factors and with a few exceptions are frequent in cancers that arise from tissues with a low rate of self-renewal^13,14,27^. The *TERT* promoter mutations affect the process of tumorigenesis through rejuvenation of telomerase through an increased *TERT* transcription^13,14^. Like in melanoma, *TERT* promoter mutations in BCC tumors are highly prevalent^28,29^. We observed that *TERT* promoter with mutations in 59% of tumors in consistent with earlier reports was the most frequently altered loci^28,29^. Of the two main *TERT* promoter mutations, -124 C>T and -146 C>T, the former, with the exception of skin neoplasms, is overwhelmingly predominant in most cancers^13^. However, in melanoma and keratinocyte cancers, the -146 C>T is the most frequent *TERT* promoter mutation as in this study. Skin cancers are also characterized by the presence of CC>TT tandem mutations at - 124/-125 and -138/-139 bp positions that lead to similar CCGGAA consensus binding site for ETS transcription factors^30^. The pattern of *TERT* promoter mutations in skin cancers is indicative of an UV etiology in the genesis of those alterations. The -138/-139 CC>TT tandem mutation in melanoma was shown to be associated with the worst melanoma-specific patient survival^31^. The altered site due to the -146 C>T mutation specifically involves non-canonical NF-kB signaling with cooperative binding between p52/RelB and ETS1^32,33^. In general sites created by *TERT* promoter mutations involve binding by multimeric GABPA/GABPB1 complex at ETS transcription factor leading to an epigenetic switch from inactive to active histone mark and pol II recruitment^15^. As reported in previous studies, we detected increased transcription of the reverse transcriptase subunit in BCC tumors with than without the *TERT* promoter mutations^34,35^.

We also investigated THOR region where methylation of CpG sites is generally associated with increased *TERT* expression^16,36^. However, our data show that increased *TERT* expression in BCC tumors can be primarily attributed to the promoter mutations as shown in adult gliomas^37^. Telomeres in tumors with the *TERT* promoter mutations are usually shorter than in tumors without those mutations as observed in melanoma and gliomas^35,38^. However, we did not observe a statistically significant difference in telomere length in BCC tumors.

Noncoding mutations within the *DPH3* promoter were detected in 38% of the lesions. The effect of the mutations, located within the proximity to an Ets binding motif, has been rather ambiguous^17^. The presence of mutations exclusively at dipyrimidinic sites coupled with typical CC>TT tandem alterations in the *DPH3* promoter mutations was indicative of an UV etiology. In the absence of a definitively ascribed functionality, the *DPH3* promoter mutations as being mere passenger events cannot be ruled out^39^. However, we observed that both *TERT* promoter and *DPH3* mutations associated with an increased risk of cutaneous neoplasms. *DPH3* encodes a short peptide involved in electron transfer during the synthesis of eukaryotic diphthamide and forms a complex with Kti13, which is involved in both tRNA and translational elongation factor 2 (EF2) modifications^40,41^. Overexpression of *DPH3* has been shown to promote migratory ability of murine melanoma and downregulation of its expression was shown to inhibit cellular invasion and metastasis in vivo42,43.

In conclusion, we found noncoding mutations within the *TERT* and *DPH3* promoter at high frequency in BCC tumors in addition to frequent alterations in *PTCH1* and *TP53* genes that impair protein functions. Interestingly, the alterations in *PTCH1, TP53* and *DPH3* promoter occurred more frequently in tumors with *TERT* promoter mutations. It is likely that the increased cellular proliferation following the activation of Hedgehog signaling or elimination of checkpoints due to p53 loss would require telomere buttressing due to increased cellular proliferation, which is probably attained through telomerase rejuvenation via *TERT* promoter mutations.

## Material and Methods

### Patients and tissues

Fresh-frozen tissues from BCC tumors from 191 patients and apparently normal appearing skin surrounding tumor tissues from 115 patients were included in this study. Seventy-one (37.2%) nodular, 42 (22.0%) superficial, and 4 (2.1%) adenoid tumors were grouped as low risk. Twenty-four (12.6%) infiltrative, 7 (3.7%) were morphoeic, 6 (3.1%) micronodular, and one metatypical type were categorized as high risk. For 36 (18.9%) tumors histological data were not available. BCC lesions were retrieved from the Biobank of the Instituto Valenciano de Oncología in Valencia, Spain and collected at the Department of Dermatology of the University of L’Aquila, Italy. Approval for this study was obtained from the Local Ethics Committee. A written informed consent was signed by all study participants.

Demographic and clinical characteristics of patients including age, sex, skin type, sun exposure, nevus count, solar lentigines, other skin cancers, immunosuppression, anatomical site and histological subtype of tumors were registered through a standardized questionnaire, skin examination and medical records **(Supplementary Table 1)**. During surgical excision of the lesions, a 4 mm intra-tumoral punch biopsy specimen was obtained, snap-frozen in liquid nitrogen and kept at -80°C until nucleic acid extraction. The remaining tissue was formalin-fixed and paraffin-embedded (FFPE) for conventional histopathology.

### Mutational Analysis by Sanger Sequencing

DNA and total RNA were extracted from fresh-frozen tissues using the QIAGEN AllPrep DNA/RNA/miRNA Universal Kit (QIAGEN, Hilden, Germany) according to the manufacturer’s instruction. Mutations at different loci were screened using PCR and Sanger sequencing using primers and standard conditions (**Supplementary Table 6**). Sequencing data were analyzed using Geneious Pro 5.6.5 software and sequences from the NCBI gene database were used as references, *PTCH1* (chr9: 98,205,262-98,279,339 hg 19 coordinates), *TP53* (chr17: 7,565,097-7,590,856 hg19 coordinates), *TERT* promoter (chr5: 1,295,071-1,295,521, hg 19 coordinates), and *DPH3* promoter (chr3: 16,306,256-16,306,755, hg19 coordinates),. The point mutations in the *PTCH1* and *TP53* genes were annotated by using web-based tool Mutalyzer^44^. The sequence topology for the *PTCH1* gene was generated with PROTTER^45^. Mutations with intron/exon boundaries were analyzed for the effect on splicing using maximum entropy model^46^. For splice site analysis on the 5’ end, three nucleotides from exon and 6 nucleotides from the following intron were included in the model; for 3’ splice site, 20 nucleotides from the intron and three nucleotides from the preceding exon were included in the model.

### Multiplex ligation-dependent probe amplification (MLPA)

MLPA method with specific probes was used (SALSA MLPA P067-B2 *PTCH1* probemix, MRC-Holland, Amsterdam, The Netherlands) to detect deletions/duplications in the *PTCH1* gene. The results were cross-validated by Sanger sequencing and concordant results were confirmed by both methods.

### Measurement of TERT promoter methylation

A 213 bp genomic region within the *TERT* promoter, from -560 to -774 (chr5:1,295,665-1,295,878; hg19) from ATG start site was screened for CpG sites in-silico using MethPrimer^47^. The selected region included 14 CpG sites and primers were designed to amplify both methylated and unmethylated sequence. PCR product was either sequenced directly or after cloning into T-overhang vector. The sequence data files were further analyzed for CpG methylation status using a web-based bisulfite sequencing analysis tool called QUMA (QUantification tool for *Methylation* Analysis) under default settings^48^.

### Measurement of TERT mRNA expression by quantitative real-time PCR (qRT-PCR)

For measurement of gene expression, reverse transcription reaction was performed using approximately 1.0 µg RNA and random hexamer primers using a cDNA synthesis kit (Thermo Scientific, Waltham, USA). The real-time PCR was carried out in triplicates on a 384-well layout using primers specific for *TERT* (**Supplementary Table 6**) and primers for the *GUSB* gene (Qiagen). Difference in gene expression levels were calculated following the ΔΔC_T_ method; GUSB expression was used as an internal reference (ΔC_T_) and difference in expression levels was calculated between BCCs and matched tumor-surrounding skin (ΔΔC_T_) followed by performing a log_2_ transformation.

### Measurement of telomere length

Relative telomere length in tumor DNA was measured using the monochrome multiplex PCR assay as described previously including minor modifications ^49,50^. The standard curve was used to quantify the telomere (T) and albumin genes (S) based on the respective average Ct values obtained in triplicate. The relative telomere length was expressed as the ratio between T/S. Inter-assay and intra-assay variation were determined by duplicating the reference DNA for all the dilutions in all the assays performed.

### Statistics

The associations between mutations in *PTCH1, TP53, TERT* promoter, *DPH3* promoter and patient age at diagnosis, sex, phototype, nevus count, solar lentigos, history of cutaneous neoplasms, exposure to sun, immunosuppressivs treatment and histology were determined by χ^2^-tests and size of the effect determined by odds ratio (OR) and 95% confidence intervals (CIs) in logistic regression model. Multivariate logistic regression was also carried and included statistically significant variables from univariate analysis. Box plots were drawn to for *TERT* expression and telomere length in tumors with and without *TERT* promoter mutations and differences analyzed by two-tailed t-test.

## Acknowledgement

The study was supported by a TRANSCAN (JTC2013) award though German Ministry of Education and Research (BMBF) under grant number 01KT15511. Patient material for the study was retrieved from the biobank of the Instituto Valenciano de Oncologia.

## Figure Legends

**Supplementary Figure 1.** Schematic representation of the *PTCH1* gene using PROTTER; all mutations identified in the BCC tumors are highlighted manually.

**Supplementary Figure 2.** Methylation pattern at 14 CpG sites at the *TERT* promoter (chr5:1,295,665-1,295,878; hg19 coordinates) determined by bisulfite modification, cloning, sequencing and analysis using QUMA. The upper panel represents methylation levels in 10 tumors without the lower panel represents methylation levels in 10 tumors with the *TERT* promoter mutations. Each circle represents a CpG site with corresponding genomic location at the top. Black colored region of the circles denotes methylated (percent) and the light striped region denotes unmethylated (percent). The statistical difference in methylation patterns is shown underneath each CpG site.

**Supplementary Figure 3.** Relative *TERT* gene expressions in BCCs with and without *TERT* promoter mutations based on *TERT* methylation status: Experiments were carried out in triplicates and box plots represent mean ± standard error of means; *P*-value was determined by *t*-test.

**Supplementary Figure 4. Telomere lengths in BCC tumors and surrounding skin**: Relative telomere length in surrounding skin, in BCC without and with *TERT* promoter mutations. Experiments were carried out in triplicates and box plot represent mean ± S.E. *P-*value was determined by *t*-test.

## References

1. Epstein EH. Basal cell carcinomas: attack of the hedgehog. Nature reviews Cancer 2008, 8(10): 743–754.

2. Peris K, Fargnoli MC, Garbe C, Kaufmann R, Bastholt L, Seguin NB, et al. Diagnosis and treatment of basal cell carcinoma: European consensus-based interdisciplinary guidelines. European journal of cancer (Oxford, England : 1990) 2019, 118: 10–34.

3. Verkouteren JAC, Ramdas KHR, Wakkee M, Nijsten T. Epidemiology of basal cell carcinoma: scholarly review. The British journal of dermatology 2017, 177(2): 359–372.

4. Youssef KK, Van Keymeulen A, Lapouge G, Beck B, Michaux C, Achouri Y, et al. Identification of the cell lineage at the origin of basal cell carcinoma. Nature cell biology 2010, 12(3): 299–305.

5. Peterson SC, Eberl M, Vagnozzi AN, Belkadi A, Veniaminova NA, Verhaegen ME, et al. Basal cell carcinoma preferentially arises from stem cells within hair follicle and mechanosensory niches. Cell stem cell 2015, 16(4): 400–412.

6. Pellegrini C, Maturo MG, Di Nardo L, Ciciarelli V, Gutierrez Garcia-Rodrigo C, Fargnoli MC. Understanding the Molecular Genetics of Basal Cell Carcinoma. International journal of molecular sciences 2017, 18(11).

7. Kasper M, Jaks V, Hohl D, Toftgard R. Basal cell carcinoma - molecular biology and potential new therapies. The Journal of clinical investigation 2012, 122(2): 455–463.

8. Bonilla X, Parmentier L, King B, Bezrukov F, Kaya G, Zoete V, et al. Genomic analysis identifies new drivers and progression pathways in skin basal cell carcinoma. Nature genetics 2016, 48(4): 398–406.

9. Sharpe HJ, Pau G, Dijkgraaf GJ, Basset-Seguin N, Modrusan Z, Januario T, et al. Genomic analysis of smoothened inhibitor resistance in basal cell carcinoma. Cancer cell 2015, 27(3): 327–341.

10. Atwood SX, Sarin KY, Whitson RJ, Li JR, Kim G, Rezaee M, et al. Smoothened variants explain the majority of drug resistance in basal cell carcinoma. Cancer cell 2015, 27(3): 342–353.

11. Jayaraman SS, Rayhan DJ, Hazany S, Kolodney MS. Mutational landscape of basal cell carcinomas by whole-exome sequencing. The Journal of investigative dermatology 2014, 134(1): 213–220.

12. Jayaraman SS, Rayhan DJ, Hazany S, Kolodney MS. Mutational landscape of basal cell carcinomas by whole-exome sequencing. The Journal of investigative dermatology 2014, 134(1): 213–220.

13. Heidenreich B, Kumar R. TERT promoter mutations in telomere biology. Mutat Res 2017, 771: 15–31.

14. Horn S, Figl A, Rachakonda PS, Fischer C, Sucker A, Gast A, et al. TERT promoter mutations in familial and sporadic melanoma. Science (New York, NY) 2013, 339(6122): 959–961.

15. Stern JL, Theodorescu D, Vogelstein B, Papadopoulos N, Cech TR. Mutation of the TERT promoter, switch to active chromatin, and monoallelic TERT expression in multiple cancers. Genes & development 2015, 29(21): 2219–2224.

16. Stern JL, Paucek RD, Huang FW, Ghandi M, Nwumeh R, Costello JC, et al. Allele-Specific DNA Methylation and Its Interplay with Repressive Histone Marks at Promoter-Mutant TERT Genes. Cell reports 2017, 21(13): 3700–3707.

17. Denisova E, Heidenreich B, Nagore E, Rachakonda PS, Hosen I, Akrap I, et al. Frequent DPH3 promoter mutations in skin cancers. Oncotarget 2015, 6(34): 35922–35930.

18. Ingham PW, Nakano Y, Seger C. Mechanisms and functions of Hedgehog signalling across the metazoa. Nature reviews Genetics 2011, 12(6): 393–406.

19. Lindstrom E, Shimokawa T, Toftgard R, Zaphiropoulos PG. PTCH mutations: distribution and analyses. Human mutation 2006, 27(3): 215–219.

20. Gailani MR, Leffell DJ, Ziegler A, Gross EG, Brash DE, Bale AE. Relationship between sunlight exposure and a key genetic alteration in basal cell carcinoma. Journal of the National Cancer Institute 1996, 88(6): 349–354.

21. Cui R, Widlund HR, Feige E, Lin JY, Wilensky DL, Igras VE, et al. Central role of p53 in the suntan response and pathologic hyperpigmentation. Cell 2007, 128(5): 853–864.

22. Levine AJ, Oren M. The first 30 years of p53: growing ever more complex. Nature reviews Cancer 2009, 9(10): 749–758.

23. Boettcher S, Miller PG, Sharma R, McConkey M, Leventhal M, Krivtsov AV, et al. A dominant-negative effect drives selection of TP53 missense mutations in myeloid malignancies. Science (New York, NY) 2019, 365(6453): 599–604.

24. Bolshakov S, Walker CM, Strom SS, Selvan MS, Clayman GL, El-Naggar A, et al. p53 mutations in human aggressive and nonaggressive basal and squamous cell carcinomas. Clinical cancer research : an official journal of the American Association for Cancer Research 2003, 9(1): 228–234.

25. Reifenberger J, Wolter M, Knobbe CB, Kohler B, Schonicke A, Scharwachter C, et al. Somatic mutations in the PTCH, SMOH, SUFUH and TP53 genes in sporadic basal cell carcinomas. The British journal of dermatology 2005, 152(1): 43–51.

26. Fell GL, Robinson KC, Mao J, Woolf CJ, Fisher DE. Skin beta-endorphin mediates addiction to UV light. Cell 2014, 157(7): 1527–1534.

27. Heidenreich B, Kumar R. Altered TERT promoter and other genomic regulatory elements: occurrence and impact. International journal of cancer 2017, 141(5): 867–876.

28. Griewank KG, Murali R, Schilling B, Schimming T, Moller I, Moll I, et al. TERT Promoter Mutations Are Frequent in Cutaneous Basal Cell Carcinoma and Squamous Cell Carcinoma. PloS one 2013, 8(11): e80354.

29. Populo H, Boaventura P, Vinagre J, Batista R, Mendes A, Caldas R, et al. TERT promoter mutations in skin cancer: the effects of sun exposure and X-irradiation. The Journal of investigative dermatology 2014, 134(8): 2251–2257.

30. Nagore E, Heidenreich B, Rachakonda S, Garcia-Casado Z, Requena C, Soriano V, et al. TERT promoter mutations in melanoma survival. International journal of cancer 2016, 139(1): 75–84.

31. Andres-Lencina JJ, Rachakonda S, Garcia-Casado Z, Srinivas N, Skorokhod A, Requena C, et al. TERT promoter mutation subtypes and survival in stage I and II melanoma patients. International journal of cancer 2019, 144(5): 1027–1036.

32. Li Y, Zhou QL, Sun W, Chandrasekharan P, Cheng HS, Ying Z, et al. Non-canonical NF-kappaB signalling and ETS1/2 cooperatively drive C250T mutant TERT promoter activation. Nature cell biology 2015, 17(10): 1327–1338.

33. Xu X, Li Y, Bharath SR, Ozturk MB, Bowler MW, Loo BZL, et al. Structural basis for reactivating the mutant TERT promoter by cooperative binding of p52 and ETS1. Nature communications 2018, 9(1): 3183.

34. Heidenreich B, Nagore E, Rachakonda PS, Garcia-Casado Z, Requena C, Traves V, et al. Telomerase reverse transcriptase promoter mutations in primary cutaneous melanoma. Nature communications 2014, 5: 3401.

35. Heidenreich B, Rachakonda PS, Hosen I, Volz F, Hemminki K, Weyerbrock A, et al. TERT promoter mutations and telomere length in adult malignant gliomas and recurrences. Oncotarget 2015, 6(12): 10617–10633.

36. Lee DD, Leao R, Komosa M, Gallo M, Zhang CH, Lipman T, et al. DNA hypermethylation within TERT promoter upregulates TERT expression in cancer. The Journal of clinical investigation 2019, 129(1): 223–229.

37. Arita H, Narita Y, Takami H, Fukushima S, Matsushita Y, Yoshida A, et al. TERT promoter mutations rather than methylation are the main mechanism for TERT upregulation in adult gliomas. Acta neuropathologica 2013, 126(6): 939–941.

38. Rachakonda S, Kong H, Srinivas N, Garcia-Casado Z, Requena C, Fallah M, et al. Telomere length, telomerase reverse transcriptase promoter mutations, and melanoma risk. Genes, chromosomes & cancer 2018, 57(11): 564–572.

39. Weinhold N, Jacobsen A, Schultz N, Sander C, Lee W. Genome-wide analysis of noncoding regulatory mutations in cancer. Nature genetics 2014, 46(11): 1160–1165.

40. Dong M, Su X, Dzikovski B, Dando EE, Zhu X, Du J, et al. Dph3 is an electron donor for Dph1-Dph2 in the first step of eukaryotic diphthamide biosynthesis. J Am Chem Soc 2014, 136(5): 1754–1757.

41. Glatt S, Zabel R, Vonkova I, Kumar A, Netz DJ, Pierik AJ, et al. Structure of the Kti11/Kti13 heterodimer and its double role in modifications of tRNA and eukaryotic elongation factor 2. Structure (London, England : 1993) 2015, 23(1): 149–160.

42. Wang L, Shi Y, Ju P, Liu R, Yeo SP, Xia Y, et al. Silencing of diphthamide synthesis 3 (Dph3) reduces metastasis of murine melanoma. PloS one 2012, 7(11): e49988.

43. Liu S, Wiggins JF, Sreenath T, Kulkarni AB, Ward JM, Leppla SH. Dph3, a small protein required for diphthamide biosynthesis, is essential in mouse development. Molecular and cellular biology 2006, 26(10): 3835–3841.

44. Wildeman M, van Ophuizen E, den Dunnen JT, Taschner PE. Improving sequence variant descriptions in mutation databases and literature using the Mutalyzer sequence variation nomenclature checker. Human mutation 2008, 29(1): 6–13.

45. Omasits U, Ahrens CH, Muller S, Wollscheid B. Protter: interactive protein feature visualization and integration with experimental proteomic data. Bioinformatics (Oxford, England) 2014, 30(6): 884–886.

46. Yeo G, Burge CB. Maximum entropy modeling of short sequence motifs with applications to RNA splicing signals. Journal of computational biology : a journal of computational molecular cell biology 2004, 11(2-3): 377–394.

47. Li LC, Dahiya R. MethPrimer: designing primers for methylation PCRs. Bioinformatics (Oxford, England) 2002, 18(11): 1427–1431.

48. Kumaki Y, Oda M, Okano M. QUMA: quantification tool for methylation analysis. Nucleic acids research 2008, 36(Web Server issue): W170–175.

49. Cawthon RM. Telomere length measurement by a novel monochrome multiplex quantitative PCR method. Nucleic acids research 2009, 37(3).

50. Shen M, Cawthon R, Rothman N, Weinstein SJ, Virtamo J, Hosgood HD, et al. A prospective study of telomere length measured by monochrome multiplex quantitative PCR and risk of lung cancer. Lung Cancer 2011, 73(2): 133–137.

