## Supplementary data for "Coding and noncoding somatic mutations in basal cell carcinoma"

Supplementary Table 1

| Tumor ID | Age | Sex | TERT promoter | DPH3 promoter | TP53 | PTCH1 | MLPA (PTCH1) | Inmunosupression | Sunexposure at work pla | Fitzpatr | Nevus | Lentigos | History of c |
|---|---|---|---|---|---|---|---|---|---|---|---|---|---|
| 1 | 74 | M | -146C>T | wt | c.529C>T | c.1249C>T | LOH | no | none | 3 | <50 | abundant | no |
| 2 | 72 | M | -146C>T | wt | wt | c.3449C>T | LOH | no | none | 3 | <50 | scarce | yes |
| 3 | 93 | F | wt | wt | wt | wt | | no | none | 3 | <50 | scarce | yes |
| 4 | 68 | F | -146C>T | -8 9CC>TT | c.880G>T | c.1725 1726CC>TT | LOH | no | none | 2 | <50 | scarce | yes |
| 5 | 75 | F | wt | wt | wt | wt | | no | <20 years | 1 | <50 | abundant | no |
| 6 | 32 | F | wt | wt | wt | c.1216-1G>A | | no | <20 years | 2 | <50 | none | no |
| 7 | 75 | F | wt | wt | c.637C>T | wt | | no | >20 years | 3 | <50 | abundant | no |
| 8 | 47 | F | wt | wt | wt | c.1347G>A; c.738 746del9 | LOH | no | none | 3 | <50 | scarce | yes |
| 9 | 90 | F | -146C>T | wt | wt | wt | | no | >20 years | 3 | <50 | abundant | no |
| 10 | 85 | M | wt | wt | wt | c.4321C>T; c.2307 2308CC>TT | LOH | no | <20 years | 3 | <50 | scarce | no |
| 11 | 67 | F | wt | wt | c.637C>T | c.3449C>T | | no | <20 years | 2 | <50 | abundant | no |
| 12 | 70 | F | wt | wt | wt | wt | | no | <20 years | 2 | <50 | abundant | no |
| 13 | 51 | F | -124C>T | wt | c.637C>T; c.733 734GG>AA | wt | LOH | no | >20 years | 2 | <50 | abundant | no |
| 14 | 61 | F | -146C>T | wt | wt | wt | | no | >20 years | 2 | <50 | scarce | yes |
| 15 | 67 | M | -101C>T | -8C>T | c.529 530del18; c.725G>T | c.1728G>A;c.1728+1G>A | LOH | no | >20 years | 2 | <50 | abundant | no |
| 16 | 77 | M | -138 139CC>TT | wt | c.772G>A; c.839G>A | wt | | no | none | 2 | <50 | scarce | no |
| 17 | 45 | M | wt | -8C>T | wt | wt | | no | none | 3 | <50 | abundant | no |
| 18 | 57 | M | -146C>T | wt | c.535C>T; c.722C>T | c.4024C>G; c.1380 1381GG>AA; c.3355delC | | no | >20 years | 2 | >50 | abundant | yes |
| 19 | 83 | M | wt | wt | c.580C>T; c.672G>A;c.672+1G>A | wt | | no | none | 2 | <50 | scarce | no |
| 20 | 75 | F | wt | -8 9CC>TT | wt | wt | | no | none | 3 | <50 | abundant | yes |
| 21 | 75 | F | wt | wt | wt | c.1504-8T>C | | no | none | 3 | <50 | scarce | yes |
| 22 | 75 | F | -146C>T | -8C>T | c.534 535CC>TT | c.1504-8T>C | | no | none | 3 | <50 | scarce | yes |
| 23 | 40 | M | wt | wt | wt | c.3487G>A | | no | <20 years | 3 | >50 | abundant | yes |
| 24 | 54 | F | -146C>T | wt | wt | c.2189 2190GG>AA | | no | none | 2 | <50 | abundant | no |
| 25 | 73 | M | -101C>T | -8 9CC>TT | wt | wt | | no | <20 years | 3 | <50 | abundant | no |
| 26 | 46 | F | -146C>T | -12C>T | wt | c.3168+3C>G | | no | none | 2 | <50 | scarce | no |
| 27 | 81 | M | -146C>T | -8C>T | wt | c.1249C>T | | no | <20 years | 2 | <50 | abundant | no |
| 28 | 68 | M | wt | wt | wt | wt | | no | none | 2 | <50 | abundant | no |
| 29 | 83 | M | -146C>T | . | wt | wt | | no | none | 3 | <50 | abundant | yes |
| 30 | 68 | M | -146C>T | -8C>T | c.813 814GG>AA; c.991C>T | c.3054G>A | | no | none | 3 | <50 | abundant | no |
| 31 | 69 | F | wt | wt | wt | c.2552 2553GG>AA | | no | >20 years | 2 | <50 | abundant | no |
| 32 | na | 1 | -138 139CC>TT | wt | c.748 749CC>TT; c.856G>A | c.654G>A;c.654+1G>A | LOH | na | none | na | na | na | na |
| 33 | 81 | F | wt | -8C>T; -12 C>T | c.535C>T; c.892G>T | c.1414delGinsTT | LOH | no | none | 4 | <50 | abundant | no |
| 34 | 75 | F | -101C>T | wt | c.531 532CC>TG | wt | | yes | <20 years | 2 | <50 | scarce | no |
| 35 | 57 | M | wt | wt | wt | c.882C>T | | no | <20 years | 2 | <50 | scarce | yes |
| 36 | 67 | M | wt | wt | wt | wt | | no | <20 years | 3 | <50 | abundant | no |
| 37 | 67 | M | wt | wt | wt | wt | | no | <20 years | 3 | <50 | abundant | no |
| 38 | 67 | M | -125C>T | wt | wt | wt | | no | <20 years | 3 | <50 | abundant | no |
| 39 | 67 | M | wt | wt | wt | wt | | no | <20 years | 3 | <50 | abundant | no |
| 40 | 47 | M | wt | wt | wt | wt | | na | none | 3 | na | scarce | no |
| 41 | 38 | F | wt | wt | wt | wt | | yes | >20 years | 3 | <50 | none | no |
| 42 | 47 | M | wt | -9C>T | wt | c.1436delT | LOH | no | >20 years | 3 | >50 | scarce | no |
| 43 | 42 | F | wt | -8C>T | wt | c.945+1C>T | LOH | no | >20 years | 3 | <50 | abundant | no |
| 44 | 69 | F | wt | wt | wt | c.672C>A | LOH | no | <20 years | 2 | <50 | scarce | no |
| 45 | 78 | M | -124 125CC>TT | -8C>T | wt | c.2561G>A; c.2001delGTACinsTTA | | no | <20 years | 2 | <50 | abundant | no |
| 46 | 68 | F | -126C>T | -8C>T | c.586C>T | c.4080C>T; c.945+2T>A | | na | none | na | na | na | no |
| 47 | 77 | M | wt | wt | wt | wt | | yes | <20 years | 3 | <50 | abundant | no |
| 48 | 82 | M | -146C>T | wt | wt | c.1119C>T | | no | none | 3 | <50 | abundant | yes |
| 49 | 50 | F | wt | wt | wt | c.3257T>C | LOH | no | <20 years | 3 | <50 | abundant | no |
| 50 | 58 | M | -146C>T | wt | c.596G>A | wt | | no | none | 2 | <50 | abundant | no |
| 51 | 50 | M | wt | wt | c.832C>T | c.3574C>T | | no | none | 3 | <50 | scarce | no |
| 52 | 77 | M | -146C>T | -8 9CC>TT | wt | wt | | no | >20 years | 2 | na | na | yes |
| 53 | 71 | M | wt | wt | wt | c.3237 3262dup21 | | no | none | 3 | na | na | no |
| 54 | 70 | M | wt | -8_9CC>TT | c.742C>T; c.832C>T | wt | LOH | no | <20 years | 3 | na | na | no |
| 55 | 79 | 2 | -124C>T | wt | wt | wt | | no | <20 years | 3 | <50 | abundant | na |
| 56 | 67 | 2 | wt | wt | wt | wt | | yes | na | 3 | <50 | scarce | yes |
| 57 | . | 2 | wt | wt | wt | wt | | na | na | na | na | na | na |
| 58 | 90 | 1 | wt | wt | wt | wt | | no | na | 2 | <50 | scarce | yes |
| 59 | 55 | 1 | wt | wt | wt | wt | | no | <20 years | 3 | >50 | scarce | yes |
| 60 | 76 | 1 | wt | wt | wt | wt | | no | none | 4 | <50 | scarce | no |
| 61 | 84 | 2 | -146C>T | -9C>T | wt | wt | LOH | no | na | 3 | na | abundant | yes |
| 62 | 82 | 1 | -124 125CC>TT | wt | wt | c.4141G>A | | na | na | na | na | na | no |
| 63 | 69 | 1 | wt | -8C>T | c.374G>A; c.458C>T; c.859G>A | c.2110 2111delAG | LOH | no | none | 3 | >50 | scarce | yes |
| 64 | 65 | 1 | wt | wt | c.528C>G; c.818G>A | c.2062C>T | LOH | no | none | 3 | >50 | abundant | no |
| 65 | 90 | 1 | -146C>T | -8C>T | wt | c.2848C>T | Focal deletion (exons 4-5 | no | na | 2 | <50 | scarce | yes |
| 66 | 63 | 1 | -146C>T | wt | c.530 531CC>TT; c.749 750CC>TT; c.722C>T; c.375-23delAG | c.1725 1726CC>GT; c.2306 2314insC | LOH | no | >20 years | 2 | >50 | abundant | yes |
| 67 | 68 | 2 | -146C>T | -9C>T | c.413C>T; c.586C>T | c.1804C>T | | no | none | 2 | <50 | none | na |
| 68 | 83 | 2 | wt | wt | wt | wt | | no | na | 1 | <50 | none | yes |
| 69 | 65 | 2 | -146C>T, -101C>T | wt | wt | c.591G>A; c.1216-6C>A; c.2703+2T>G; c.3921 3922insC | | na | na | 2 | >50 | na | na |
| 70 | 76 | 1 | -124C>T | -9C>A | c.637C>T; c.585 586CC>TT; c.839G>A | c.2689A>G | | no | >20 years | 4 | <50 | none | yes |
| 71 | 82 | 2 | wt | wt | wt | c.945+1C>T | LOH | no | none | 2 | <50 | abundant | na |
| 72 | 80 | 1 | -146C>T | wt | wt | c.3168+2T>A | | no | na | 2 | <50 | abundant | yes |
| 73 | 84 | 1 | wt | -12C>T | wt | c.3182delCGCinsTG | | no | >20 years | 2 | <50 | scarce | yes |
| 74 | . | 1 | wt | wt | wt | wt | | na | na | na | na | na | na |
| 75 | .80 | 2 | -146C>T | wt | wt | c.569 571delTAT | | na | none | 2 | na | na | no |
| 76 | 83 | 1 | wt | -8C>T | c.722C>T | wt | | na | na | 2 | na | na | yes |
| 77 | 64 | 1 | wt | wt | wt | wt | | no | none | 2 | <50 | scarce | yes |
| 78 | 52 | 1 | -124C>T | -8C>T | wt | c.4321C>T | | no | none | 3 | <50 | scarce | yes |
| 79 | 86 | 1 | wt | wt | wt | wt | | no | <20 years | 4 | <50 | abundant | yes |
| 80 | 80 | 1 | -124C>T | -8C>T | wt | c.2599 2604insAAATC | | na | >20 years | 4 | <50 | scarce | yes |
| 81 | 80 | 1 | -146C>T | -8 9CC>TT | c.464C>T; c.596C>T; c.586C>T; c.722C>T | c.1526G>A | | no | >20 years | 3 | <50 | scarce | na |
| 82 | 84 | 2 | wt | wt | wt | wt | | yes | <20 years | 4 | <50 | scarce | na |
| 83 | 87 | 2 | -124C>T | wt | wt | wt | | no | none | 3 | na | abundant | na |
| 84 | 39 | 2 | -146C>T | -8 9CC>TT | c.459delC; c.836G>A; c.832C>T; c.948 949GG>AA | c.3053G>A | LOH | no | none | 4 | >50 | scarce | na |
| 85 | 76 | 1 | -146C>T | -9C>T | c.530C>T; c.454 455CC>TT; c.672+2A>T | c.619G>A | LOH | no | na | 2 | <50 | scarce | yes |
| 86 | . | 2 | -124 125CC>TT | -8C>T | wt | c.834G>A | | na | na | na | na | na | na |
| 87 | 74 | 1 | -138 139CC>TT | wt | wt | wt | | no | none | 3 | <50 | abundant | no |
| 88 | 71 | 1 | wt | -8C>T | wt | wt | | na | na | na | na | na | no |
| 89 | 33 | 1 | wt | wt | wt | wt | | no | none | 3 | >50 | abundant | yes |
| 90 | . | 1 | -146C>T | wt | c.839G>T | c.855 865del 11 | | na | na | na | na | na | na |
| 91 | 69 | 1 | -124C>T | wt | c.707A>G | c.1161G>A | Focal deletion (exons 18- | no | none | 3 | >50 | abundant | yes |
| 92 | 51 | 1 | -146C>T, -101C>T | -8C>T | wt | c.539 545delACTCGGC; c.2887+4del AGTA | | no | none | 3 | <50 | scarce | no |
| 93 | 73 | 2 | -146C>T | -8C>T | wt | c.1380G>A; c.3027C>T | | no | >20 years | 4 | <50 | scarce | na |
| 94 | 87 | 1 | -138 139CC>TT | -8C>T | c.574C>T; c.733 734GG>TT | c.4213G>A; c.3168G>A;c.3168-1G>A | | na | na | na | na | na | na |
| 95 | 47 | 1 | -146C>T | wt | c.741 742CC>TT | c.3277G>A | LOH | no | none | 2 | >50 | scarce | yes |
| 96 | 72 | 2 | -146C>T, -149C>T | wt | wt | wt | | no | none | 4 | <50 | scarce | no |
| 97 | 83 | 2 | -146C>T | wt | wt | wt | | no | none | na | <50 | na | no |
| 98 | 80 | 2 | wt | wt | c.406C>T | c.1703C>T; c.1340delA | LOH | no | none | 3 | >50 | abundant | yes |
| 99 | 77 | 1 | -146C>T | wt | wt | c.2770 2788del21 | | no | none | 2 | <50 | none | no |
| 100 | 55 | 2 | wt | wt | wt | c.1306C>T | | no | none | 3 | <50 | scarce | no |
| 101 | 74 | 2 | -146C>T | -8C>T | wt | c.3587C>T; c.2921 2922insAGTT | | no | na | 2 | <50 | scarce | yes |
| 102 | 60 | 2 | -146C>T | wt | c.637C>T; c.722C>T | c.3591C>T; c.2561-1G>A | Focal deletion (From exo | no | none | 2 | <50 | abundant | yes |
| 103 | 31 | 1 | wt | wt | wt | wt | | no | none | 4 | <50 | scarce | na |
| 104 | 80 | 1 | wt | -9C>T | c.734G>A | c.2620A>T | | no | >20 years | 2 | <50 | none | yes |
| 105 | 80 | 1 | -138 139CC>TT | -8 9CC>TT | c.530 531CC>TT; c.856G>A | c.2812C>T | | no | <20 years | 1 | <50 | abundant | yes |
| 106 | 81 | 1 | -146C>T | -8 9CC>TT | c.530C>T; c.800 801GG>AA | c.3080 3081CC>TT | LOH | no | none | 1 | >50 | none | yes |
| 107 | 41 | 2 | -146C>T | wt | wt | c.2173C>T; c.1602C>T;c.1602+1C>T; c.3322 3323insA | | no | na | 2 | <50 | scarce | yes |
| 108 | 80 | 1 | -124C>T | wt | wt | wt | | no | none | 2 | <50 | scarce | yes |
| 109 | 69 | 1 | -146C>T | wt | wt | c.1136delTCGT | | no | none | 4 | <50 | none | no |
| 110 | 81 | 1 | -138 139CC>TT | wt | wt | c.1347+2C>T | LOH | no | none | 2 | <50 | abundant | yes |
| 111 | 86 | 1 | -124C>T | -8C>T | c.637C>T | c.590G>A | LOH | yes | >20 years | 3 | <50 | scarce | na |
| 112 | 50 | 1 | wt | wt | wt | wt | | no | none | 2 | <50 | abundant | na |
| 113 | 81 | 1 | wt | wt | wt | c.3662C>T | LOH | no | none | 2 | <50 | scarce | na |
| 114 | 60 | 2 | -146C>T | wt | wt | c.1067 1068insA | | no | na | 3 | <50 | abundant | yes |
| 115 | 65 | 1 | -124C>T | wt | wt | c.896C>T; c.2768T>C; c.3945C>T | LOH | no | none | 3 | >50 | abundant | yes |
| 116 | . | 1 | wt | wt | wt | wt | | na | na | na | na | na | na |
| 117 | 76 | 1 | wt | wt | wt | wt | | no | >20 years | 3 | <50 | abundant | no |
| 118 | 61 | 1 | wt | wt | wt | wt | | no | none | 1 | na | na | na |
| 119 | 76 | 1 | -146C>T | -8 9CC>TT | c.771 772GG>AA | c.1804C>T | | no | >20 years | 2 | <50 | scarce | yes |
| 120 | 67 | 1 | wt | wt | wt | wt | | no | >20 years | 3 | <50 | abundant | yes |
| 121 | 47 | 2 | wt | wt | wt | wt | | no | none | 1 | >50 | abundant | no |
| 122 | 75 | 1 | wt | wt | wt | wt | | no | >20 years | 3 | <50 | abundant | yes |
| 123 | 76 | 1 | -146C>T, -149C>T | -9C>T | c.569C>T | wt | | no | >20 years | 3 | <50 | abundant | no |
| 124 | 70 | 1 | -101C>T | -9C>T | wt | c.1804C>T | | no | <20 years | 3 | <50 | abundant | yes |
| 125 | 45 | 2 | -138 139CC>TT | wt | wt | c.740 741insC | | no | none | 1 | <50 | abundant | na |
| 126 | 72 | 1 | -146C>T | -9C>T | wt | wt | | no | none | 2 | <50 | abundant | yes |
| 127 | 68 | 1 | -124C>T | -8C>T | wt | wt | LOH | no | none | 4 | <50 | scarce | yes |
| 128 | 57 | 2 | -146C>T | -8C>T | wt | c.820C>T | | no | none | 2 | <50 | scarce | yes |
| 129 | 77 | 1 | -146C>T, -149C>T | -9C>T | wt | wt | | no | none | 2 | <50 | abundant | na |
| 130 | 39 | 2 | -146C>T | wt | wt | c.612delC | | no | none | 2 | na | na | na |
| 131 | 72 | 1 | -138 139CC>TT | wt | wt | c.3438 3449del12 | | no | >20 years | 3 | <50 | scarce | na |
| 132 | 67 | 2 | -124C>T | -8C>T | c.535C>T; c.783-4 G>A | c.590 591GG>AA | | no | none | 2 | <50 | abundant | yes |
| 133 | 77 | 1 | -146C>T | -8 9CC>TT | wt | wt | | no | none | 2 | <50 | scarce | na |
| 134 | 79 | 1 | -146C>T | wt | c.743G>A | c.2634C>T; c.585C>T | | no | none | 3 | <50 | scarce | no |
| 135 | 54 | 1 | wt | wt | wt | wt | | yes | none | 2 | na | na | no |
| 136 | 68 | 1 | -146C>T | wt | wt | wt | | no | none | 3 | <50 | none | yes |
| 137 | 95 | 2 | -146C>T | wt | wt | wt | | no | >20 years | 3 | <50 | abundant | na |
| 138 | . | 2 | -138 139CC>TT, -125C>T | -9C>A | wt | c.2999 3029dup30 | LOH | na | na | na | na | na | na |
| 139 | 72 | 1 | wt | wt | c.530C>T; c.733 734GG>TT | c.3079 3080delTG | LOH | no | na | 3 | <50 | none | na |
| 140 | 79 | 2 | -146C>T | -8C>T | wt | wt | | no | none | 2 | <50 | scarce | na |
| 141 | 64 | 1 | wt | wt | c.574C>T | wt | | no | <20 years | 2 | <50 | scarce | yes |
| 142 | 61 | 2 | -146C>T | wt | wt | wt | | no | >20 years | 4 | <50 | abundant | yes |
| 143 | 89 | 2 | -138 139CC>TT | -8C>T | wt | wt | | no | >20 years | 3 | <50 | abundant | na |
| 144 | 94 | 2 | -124C>T, -101 102CC>TT | -9C>T | wt | c.3959G>A; c.1603-1G>A | | na | na | na | na | na | yes |
| 145 | 82 | 2 | -146C>T | wt | wt | wt | | no | none | 2 | <50 | scarce | na |
| 146 | 87 | 2 | -146C>T, -126C>T | -8 9CC>TT | c.722C>T; c.844C>T | wt | LOH | na | none | 4 | na | none | na |
| 147 | 50 | 1 | wt | -8C>T | c.743 744GG>AA | c.644 647delACAT | LOH | na | na | 2 | na | na | na |
| 148 | 44 | 1 | -146C>T; -148C>T | -8C>T | c.668C>T; c.795 796GG>AA | c.2710A>T; c.3918 3919CC>TT | | no | <20 years | 4 | >50 | scarce | na |
| 149 | 60 | 1 | wt | wt | wt | wt | | no | none | 3 | <50 | scarce | na |
| 150 | 78 | 2 | -146C>T | -8C>T | c.532C>T | c.2050G>T | LOH | no | none | 3 | <50 | abundant | na |
| 151 | 79 | 1 | -101C>T, -126C>T | -8 9CC>TT; -13C>T | c.417G>C; c.380 381CC>TT | c.3268G>T | Focal deletion (exons 5-6 | no | >20 years | 3 | <50 | none | yes |
| 152 | 82 | 1 | wt | wt | wt | wt | LOH | na | none | 3 | <50 | abundant | yes |
| 153 | 82 | 1 | -146C>T | -8C>T | c.406C>T | c.2687C>T; c.757 776dup19 | | no | na | 3 | <50 | scarce | na |
| 154 | 75 | 1 | wt | wt | wt | wt | | no | <20 years | 4 | <50 | scarce | no |
| 155 | 79 | 1 | wt | wt | wt | wt | | na | none | 3 | <50 | abundant | yes |
| 156 | 79 | 1 | -146C>T | -8C>T | wt | c.576G>A | | no | none | 2 | <50 | none | no |
| 157 | 31 | 1 | -146C>T | -9C>T | wt | c.1306G>A; c.943 945delAAA | LOH | no | <20 years | 3 | <50 | scarce | yes |
| 158 | 84 | 2 | -146C>T | wt | wt | c.518 538del21 | LOH | no | >20 years | 5 | <50 | scarce | no |
| 159 | 73 | 1 | -146C>T | wt | wt | wt | | no | none | 4 | <50 | abundant | na |
| 160 | 35 | 1 | -146C>T | wt | c.530C>T; c.637C>T | c.1068 1101del34 | LOH | no | none | 2 | <50 | abundant | no |
| 161 | 45 | 1 | -138 139CC>TT | wt | wt | c.981delT; c.1350 1351insG | | no | none | 3 | <50 | scarce | na |
| 162 | 72 | 1 | wt | wt | wt | wt | | no | none | 3 | <50 | scarce | yes |
| 163 | 61 | 1 | -146C>T | -8C>T | wt | wt | | no | none | 2 | <50 | scarce | no |
| 164 | 73 | 2 | wt | -8C>T | wt | c.655-1G>A | | no | none | na | na | scarce | no |
| 165 | 60 | 2 | wt | wt | c.637C>T; c.836G>A | c.2011delC | | no | none | 3 | <50 | none | na |
| 166 | 88 | 1 | wt | wt | wt | wt | | no | none | 3 | na | scarce | na |
| 167 | 80 | 1 | -146C>T, -101C>T | wt | wt | wt | | no | >20 years | 3 | <50 | none | yes |
| 168 | 78 | 1 | -146C>T | -8 9CC>TT; -12C>T | c.586C>T; c.856G>T | wt | | yes | none | 3 | <50 | abundant | yes |
| 169 | 90 | 1 | -146C>T | wt | wt | wt | | no | >20 years | 3 | <50 | scarce | yes |
| 170 | 54 | 2 | wt | -8C>T | c.535C>T | c.2974G>T | | no | none | 3 | >50 | scarce | na |
| 171 | 46 | 2 | -146C>T | wt | wt | c.722T>A | | no | na | 4 | <50 | none | na |
| 172 | 77 | 2 | -146C>T | wt | wt | wt | | no | none | 3 | <50 | abundant | yes |
| 173 | 88 | 2 | -124C>T | wt | wt | c.2964 2965GG>TT | | no | >20 years | 2 | <50 | abundant | na |
| 174 | 83 | 1 | wt | -8 9CC>TT | c.706T>C | c.926C>T | | no | >20 years | 1 | <50 | abundant | yes |
| 175 | 70 | 1 | -138 139CC>TT | -9C>T | c.530C>T; c.380C>T | c.2908G>T; c.3306+26delTT | | no | >20 years | 3 | <50 | abundant | yes |
| 176 | 67 | 1 | -146C>T | wt | wt | c.1234delG | | no | none | 3 | <50 | scarce | no |
| 177 | 82 | 1 | -146C>T | -8C>T | wt | c.4080C>T; c.3326 3346del21 | | no | >20 years | 1 | <50 | abundant | na |
| 178 | 78 | 1 | wt | wt | wt | wt | | no | <20 years | 3 | <50 | none | yes |
| 179 | 79 | 2 | wt | -8 9CC>TT | wt | wt | | no | none | 4 | <50 | none | yes |
| 180 | 88 | 2 | wt | wt | wt | wt | | no | none | 4 | <50 | scarce | no |
| 181 | 78 | 1 | -124C>T | wt | wt | c.2350G>T | | yes | na | na | <50 | abundant | yes |
| 182 | 76 | 2 | -146C>T | wt | wt | wt | | na | none | 3 | na | na | na |
| 183 | 81 | 2 | -146C>T | wt | wt | wt | | no | none | 4 | >50 | abundant | no |
| 184 | 60 | 1 | wt | wt | wt | wt | | no | none | 3 | >50 | abundant | no |
| 185 | 78 | 1 | -146C>T | -9C>T | c.469G>T; c.782+1G>A | c.4012C>T; c.2887+1C>T | | no | >20 years | 4 | <50 | abundant | yes |
| 186 | 51 | 2 | -149C>T | -8 9CC>TT | c.947C>T | c.857 869del13 | | no | na | 2 | <50 | abundant | na |
| 187 | 74 | 2 | -146C>T | -8 9CC>TT | c.734G>A | c.3441C>T; c.517G>T | | no | none | 2 | <50 | scarce | yes |
| 188 | 83 | 2 | -146C>T | -8 9CC>TT | wt | c.3338G>A; c.746+3A>T; c.1702 1703delCC | | no | none | 2 | <50 | scarce | na |
| 189 | 82 | 1 | -146C>T | -8C>T | c.530 531CC>TT | c.3404A>G | | no | none | 2 | <50 | abundant | yes |
| 190 | 52 | 2 | -146C>T | -9C>T | c.380C>T | c.1506 1515del10 | | no | none | 2 | <50 | abundant | na |
| 191 | 73 | 2 | wt | wt | wt | wt | | no | none | 2 | <50 | scarce | no |

**Supplementary Table 2: PTCH1 mutations in BCC tumors**

| Tumor-ID | Genomic position (NC_000009.11) | Exon /Intron | CDS mutation (NM_000264.3) | Amino acid mutation (CCDS6714) | COSMIC database | MLPA |
|---|---|---|---|---|---|---|
| 1 | g.98240435 | Exon 9 | c.1249C>T | p.Q417* | p.Q417H | LOH |
| 2 | g.98215760 | Exon 20 | c.3449C>T | p.G1149G | | LOH |
| 4 | g.98238318_98238319 | Exon 12 | c.1725_1726CC>TT | p.L575L; Q576* | | LOH |
| 6 | g.98240469 | Intron 8 | c.1216-1G>A | | c.1216-1G>A | |
| 8 | g.98240337 | Exon 9 | c.1347G>A | p.M449I | p.M449I | LOH |
| | g.98244231_98244239 | Exon 5 | c.738_746del9 | p.Y247_L249del | | |
| 10 | g.98209217 | Exon 23 | c.4321C>T | p.P1441S | p.P1441S | LOH |
| | g.98229650_98229651 | Exon 15 | c.2307_2308CC>TT | p.T769T; R770* | c.2307_2308CC>TT | |
| 11 | g.98215760 | Exon 20 | c.3449C>T | p.G1149G | | |
| 13 | | | . | | | LOH |
| 15 | g.98232213_98238316 | Exon 12/ Intron 12 | c.1728G>A;c.1728-1G>A | p.Q576Q | | LOH |
| 18 | g.98209514 | Exon 23 | c.4024C>G | p.R1342G | | |
| | g.98239951_98239952 | Exon 10 | c.1380_1381GG>AA | p.W460*; D461N | | |
| | g.98215854 | Exon 20 | c.3355delC | p.L1119fs*20 | | |
| 21 | g.98239140 | Intron 10 | c.1504-1T>C | | | |
| 22 | g.98239140 | Intron 10 | c.1504-1T>C | | | |
| 23 | g.98212185 | Exon 21 | c.3487G>A | p.G1163S | p.G1163S | |
| 24 | g.98231093_98231094 | Exon 14 | c.2189_2190GG>AA | p.W760* | | |
| 26 | g.98220292 | Intron 18 | c.3168+3C>G | | | |
| 27 | g.98240435 | Exon 9 | c.1249C>T | p.Q417* | p.Q417H | |
| 30 | g.98220409 | Exon 18 | c.3054G>A | p.W1018* | p.W1018* | |
| 31 | g.98229405_98229406 | Exon 15 | c.2552_2553GG>AA | p.W851* | p.W851* | |
| 32 | g.98244322_98244416 | Exon 4/ Intron 4 | c.654G>A;c.654-1G>A | p.Q218Q | p.Q218Q | LOH |
| 33 | g.98239918 | Exon 10 | c.1414delGinsTT | p.A472fs*25 | | LOH |
| 35 | g.98242735 | Exon 6 | c.882C>T | p.R294R | | |

| | | | | | | |
|---|---|---|---|---|---|---|
| 42 | g.98239896 | Exon 10 | c.1436delT | p.L479fs*12 | | LOH |
| 43 | g.98242671 | Intron 6 | c.945+1C>T | | | LOH |
| 44 | g.98244305 | Exon 5 | c.672C>A | p.Y224* | p.Y224fs*4 | LOH |
| 45 | g.98224280 | Exon 15 | c.2561G>A | p.G854R | p.G854R | |
| | g.98231282 | Exon 14 | c.2001delGTACinsTTA | p.E667fs*2 | | |
| 46 | g.98209458 | Exon 23 | c.4080C>T | p.S1360S | p.S1360S | |
| | g.98242670 | Intron 6 | c.945+2T>A | | | |
| 48 | g.98241378 | Exon 8 | c.1119C>T | p.Y372Y | | |
| 49 | g.98218607 | Exon 19 | c.3257T>C | p.L1086P | | LOH |
| 51 | g.98211581 | Exon 22 | c.3574C>T | p.R1192C | | |
| 53 | g.98218602_98218627 | Exon 19 | c.3237_3262dup21 | p.V1081_I1087dup | | |
| 54 | | | . | | | LOH |
| 61 | | | . | | | LOH |
| 62 | g.98209397 | Exon 23 | c.4141G>A | p.V1381M | p.V1381M | |
| 63 | g.98231172_98231173 | Exon 14 | c.2110_2111delAG | p.S704fs*33 | | LOH |
| 64 | g.98231221 | Exon 14 | c.2062C>T | p.Q688* | p.Q688* | LOH |
| 65 | g.98221921 | Exon 17 | c.2848C>T | p.H950Y | | Focal deletion (exons_4-5-6-7-8) |
| 66 | g.98238318_98238319 | Exon 12 | c.1725_1726CC>GT | p.L575L; Q576* | | LOH |
| | g.98229644_98229652 | Exon 15 | c.2306_2314insG | p.T769fs | | |
| 67 | g.98232138 | Exon 13 | c.1804C>T | p.R602* | p.R602* | |
| 69 | g.98244479 | Exon 4 | c.591G>A | p.W197* | p.W197* | |
| | g.98240474 | Intron 8 | c.1216-6C>A | | c.1216-6C>A | |
| | g.98224136 | Intron 16 | c.2703+2T>G | | | |
| | g.98209616_98209617 | Exon 23 | c.3921_3922insC | p.P1307fs*17 | c.3921_3922insC | |
| 70 | g.98224152 | Exon 16 | c.2689A>G | p.I897V | | |
| 71 | g.98242671 | Intron 6 | c.945+1C>T | | | LOH |
| 72 | g.98220293 | Intron 18 | c.3168+2T>A | | | |

| | | | | | | |
|---|---|---|---|---|---|---|
| 73 | g.98218682 | Exon 19 | c.3182delCGCinsTG | p.A1061Vfs*1 | | |
| 75 | g.98247980_98247982 | Exon 3 | c.569_571delTAT | p.V190_Y191del | | |
| 78 | g.98209217 | Exon 23 | c.4321C>T | p.P1441S | p.P1441S | |
| 80 | g.98224237_98224242 | Exon 16 | c.2599_2604insAAATC | p.K2899_I2900dup | | |
| 81 | g.98239117 | Exon 11 | c.1526G>A | p.G509D | p.G509D | |
| 84 | g.98220410 | Exon 18 | c.3053G>A | p.W1018* | p.W1018* | LOH |
| 85 | g.98244451 | Exon 4 | c.619G>A | p.G207R | | LOH |
| 86 | g.98242783 | Exon 6 | c.834G>A | p.W278* | p.W278* | |
| 90 | g.98242752_98242762 | Exon 6 | c.855_865del 11 | p.E286_H289delfs*30 | | |
| 91 | g.98241336 | Exon 8 | c.1161G>A | p.W387* | p.W387* | Focal deletion (exons_18-19) |
| 92 | g.98248006_98248012 | Exon 3 | c.539_545delACTCGGC | p.D180_A182delfs*37 | | |
| | g.98221878 | Intron 17 | c.2887+4del AGTA | | | |
| 93 | g.98239952 | Exon 10 | c.1380G>A | p.W460* | | |
| | g.98220436 | Exon 18 | c.3027C>T | p.Y1009Y | p.Y1009Y | |
| 94 | g.98209325 | Exon 23 | c.4213G>A | p.D1405N | | |
| | g.98218695_98220295 | Exon 19 /Intron 19 | c.3168G>A;c.3168-1G>A | p.V1057M | | |
| 95 | g.98218587 | Exon 19 | c.3277G>A | p.G1093R | | LOH |
| 98 | g.98238341 | Exon 12 | c.1703C>T | p.P568L | p.P568L | LOH |
| | g.98240344 | Exon 9 | c.1340delA | p.L447Tfs*9 | | |
| 99 | g.98221981_98221999 | Exon 17 | c.2770_2788del21 | p.T924Cfs*32 | | |
| 100 | g.98240378 | Exon 9 | c.1306C>T | p.D436N | | |
| 101 | g.98211568 | Exon 22 | c.3587C>T | p.P1196L | | |
| 101 | g.98220541_98220542 | Exon 18 | c.2921_2922insAGTT | p.F974* | p.F974C | |
| 102 | g.98211564 | Exon 22 | c.3591C>T | p.S1197S | p.S1197S | Focal deletion (From exon_7 to exon_25) |
| | g.98224281 | Intron 15 | c.2561-1G>A | | c.2561-1G>A | |
| 104 | g.98224221 | Exon 16 | c.2620A>T | p.K874* | | |

| | | | | | | |
|---|---|---|---|---|---|---|
| 105 | g.98221957 | Exon 17 | c.2812C>T | p.Q938* | p.Q938* | |
| 106 | g.98220382_98220383 | Exon 18 | c.3080_3081CC>TT | p.W1027* | p.W1027* | LOH |
| 107 | g.98231110 | Exon 14 | c.2173C>T | p.P725S | p.P725S | |
| | g.98238441_98239041 | Exon 11/ Intron 11 | c.1602C>T;c.1602-1C>T | p.E534N | | |
| | g.98215886_98215887 | Exon 20 | c.3322_3323insA | p.I1108Nfs*37 | | |
| 109 | g.98241361 | Exon 8 | c.1136delTCGT | p.Y379_E380delfs*52 | | |
| 110 | g.98240335 | Intron 9 | c.1347+2C>T | | | LOH |
| 111 | g.98244480 | Exon 4 | c.590G>A | p.W197* | p.W197* | LOH |
| 113 | g.98211493 | Exon 22 | c.3662C>T | p.S1221F | p.S1221F | LOH |
| 114 | g.98241429_98242251 | Intron 7 | c.1067_1068insA | | | |
| 115 | g.98242721 | Exon 6 | c.896C>T | p.P299L | | LOH |
| | g.98222001 | Exon 17 | c.2768T>C | p.L923P | p.L923P | |
| | g.98209593 | Exon 23 | c.3945C>T | p.P1315P | | |
| 119 | g.98232138 | Exon 13 | c.1804C>T | p.R602* | p.R602* | |
| 124 | g.98232138 | Exon 13 | c.1804C>T | p.R602* | p.R602* | |
| 125 | g.98244236_98244237 | Exon 5 | c.740_741insC | p.L248Pfs*4 | | |
| 127 | | | . | | | LOH |
| 128 | g.98242797 | Exon 6 | c.820C>T | p.Q274* | | |
| 130 | g.98244458 | Exon 4 | c.612delC | p.Y204* | | |
| 131 | g.98215760_98215771 | Exon 20 | c.3438_3449del12 | p.D1146_R1150del | | |
| 132 | g.98244479_98244480 | Exon 4 | c.590_591GG>AA | p.W197* | p.W197* | |
| 134 | g.98224207 | Exon 16 | c.2634C>T | p.D878D | p.D878D | |
| | g.98244485 | Intron 3 | c.585C>T | | | |
| 138 | g.98220434_98220464 | Exon 18 | c.2999_3029dup30 | p.N1000_1009dupfs | | LOH |
| 139 | g.98220383_98220384 | Exon 18 | c.3079_3080delTG | p.W1027fs*118 | | LOH |
| 144 | g.98209579 | Exon 23 | c.3959G>A | p.R1320K | p.R1320K | |
| | g.98238442 | Intron 11 | c.1603-1G>A | | c.1603-1G>A | |
| 146 | | | . | | | LOH |

| | | | | | | |
|---|---|---|---|---|---|---|
| 147 | g.98244423_98244426 | Exon 4 | c.644_647delACAT | p.Y215fs*5 | | LOH |
| 148 | g.98222059 | Exon 17 | c.2710A>T | p.K904* | p.K904>IQ | |
| | g.98209619_98209620 | Exon 23 | c.3918_3919CC>TT | p.P1306P; P1307S | | |
| 150 | g.98231233 | Exon 14 | c.2050G>T | p.E684* | p.E684* | LOH |
| 151 | g.98218596 | Exon 19 | c.3268G>T | p.V1090F | | Focal deletion (exons_5-6-7-8) |
| 152 | | | . | | | LOH |
| 153 | g.98224154 | Exon 16 | c.2687C>T | p.P896L | | |
| | g.98242841_98242860 | Exon 6 | c.757_776dup19 | p.P253_N258fs | | |
| 156 | g.98247975 | Exon 3 | c.576G>A | p.M192I | | |
| 157 | g.98240378 | Exon 9 | c.1306G>A | p.D436N | | LOH |
| | g.98242672_98242674 | Exon 6 | c.943_945delAAA | p.K315del | | |
| 158 | g.98248013_98248033 | Exon 6 | c.518_538del21 | p.E172_D180del | | LOH |
| 160 | g.98241396_98241429 | Exon 8 | c.1068_1101del34 | p.S356_M367delfs*38 | | LOH |
| 161 | g.98242337 | Exon 7 | c.981delT | p.C327fs*15 | | |
| | g.98239981_98239982 | Exon 10 | c.1350_1351insG | p.A451Gfs*46 | | |
| 164 | g.98244323 | Intron 5 | c.655-1G>A | | c.655-1G>A | |
| 165 | g.98231272 | Exon 14 | c.2011delC | p.H671Tfs*22 | | LOH |
| 168 | | | . | | | LOH |
| 170 | g.98220489 | Exon 18 | c.2974G>T | p.E992* | p.E992* | LOH |
| 171 | g.98244255 | Exon 5 | c.722T>A | p.L241* | p.L241L | |
| 173 | g.98220498_98220499 | Exon 18 | c.2964_2965GG>TT | p.V988V; E989* | | |
| 174 | g.98242691 | Exon 6 | c.926C>T | p.P309L | p.P309L | |
| 175 | g.98220555 | Exon 18 | c.2908G>T | p.E970* | p.E970* | LOH |
| | g.98218532 | Intron 19 | c.3306+26delTT | | | |
| 176 | g.98240450 | Exon 9 | c.1234delG | p.A412Hfs*27 | | |
| 177 | g.98209458 | Exon 23 | c.4080C>T | p.S1360S | p.S1360S | LOH |
| | g.98215863_98215883 | Exon 20 | c.3326_3346del21 | p.G1109_V1116del | | |

| | | | | | | |
|---|---|---|---|---|---|---|
| 181 | g.98229608 | Exon 15 | c.2350G>T | p.E784* | p.E784D | |
| 185 | g.98209526 | Exon 23 | c.4012C>T | p.R1338C | | LOH |
| | g.98221881 | Exon 17 | c.2887+1C>T | | | |
| 186 | g.98242748_98242760 | Exon 6 | c.857_869del13 | p.E286Cfs*34 | | LOH |
| 187 | g.98215768 | Exon 20 | c.3441C>T | p.F1147F | | LOH |
| | g.98248034 | Exon 3 | c.517G>T | p.E173* | p.E173V | |
| 188 | g.98215871 | Exon 20 | c.3338G>A | p.R1113H | | |
| | g.98244228 | Exon 5 | c.746+3A>T | | | |
| | g.98238341_98238342 | Exon 12 | c.1702_1703delCC | p.P568Sfs*58 | | |
| 189 | g.98215805 | Exon 16 | c.3404A>G | p.L1135P | | LOH |
| 190 | g.98239128_98239137 | Exon 11 | c.1506_1515del10 | p.V502Vfs*36 | | |

| Supplementary Table 3. Splice site mutations in PTCH1 gene | | | | | |
|---|---|---|---|---|---|
| **5' splice site mutations** | | | | | |
| | CDS location | Sequence | MAX-entropy | Markov Model | PWM |
| Mutation | c.1728+1G>A | cagATGAGC | MAXENT: 1.42 | MM: 2.12 | WMM: 2.29 |
| Wild type | c.1728+1G | cagGTGAGC | MAXENT: 9.60 | MM: 10.30 | WMM: 10.47 |
| Mutation | c.3168+3G>C | attGTCAGT | MAXENT: -0.28 | MM: -1.32 | WMM: 2.03 |
| Wild type | c.3168+3G | attGTGAGT | MAXENT: 7.35 | MM: 5.98 | WMM: 5.17 |
| Mutation | c.654+1G>A | cagATATAT | MAXENT: -0.30 | MM: -0.94 | WMM: -1.51 |
| Wild type | c.654+1G | cagGTATAT | MAXENT: 7.88 | MM: 7.24 | WMM: 6.68 |
| Mutation | c.945+1G>A | aaaATGAGT | MAXENT: 0.22 | MM: -0.40 | WMM: -0.02 |
| Wild type | c.945+1G | aaaGTGAGT | MAXENT: 8.40 | MM: 7.78 | WMM: 8.16 |
| Mutation | c.945+2T>A | aaaGAGAGT | MAXENT: 0.22 | MM: -0.40 | WMM: -0.02 |
| Wild type | c.945+2T | aaaGTGAGT | MAXENT: 8.40 | MM: 7.78 | WMM: 8.16 |
| Mutation | c.2703+2T>G | cagGGACTC | MAXENT: -0.61 | MM: -1.23 | WMM: -2.69 |
| Wild type | c.2703+2T | cagGTACTC | MAXENT: 7.04 | MM: 6.41 | WMM: 4.96 |
| Mutation | c.3168+2T>A | attGAGAGT | MAXENT: -0.83 | MM: -2.20 | WMM: -3.01 |
| Wild type | c.3168+2T | attGTGAGT | MAXENT: 7.35 | MM: 5.98 | WMM: 5.17 |
| Mutation | c.2887+3del | gaaGTAGCG | MAXENT: -3.82 | MM: 2.04 | WMM: 1.56 |
| Wild type | c.2887+3 | gaaGTAAGT | MAXENT: 9.82 | MM: 8.09 | WMM: 8.70 |
| Mutation | c.3168+1G>A | attATGAGT | MAXENT: -0.83 | MM: -2.20 | WMM: -3.01 |
| Wild type | c.3168+1G | attGTGAGT | MAXENT: 7.35 | MM: 5.98 | WMM: 5.17 |
| Mutation | c.1602+1G>A | gagATAATG | MAXENT: 0.55 | MM: -1.18 | WMM: -1.15 |
| Wild type | c.1602+1G | gagGTAATG | MAXENT: 8.73 | MM: 7.00 | WMM: 7.03 |
| Mutation | c.1347+2T>C | atgGCAACG | MAXENT: -1.71 | MM: -1.64 | WMM: -1.86 |
| Wild type | c.1347+2T | atgGTAACG | MAXENT: 6.05 | MM: 6.12 | WMM: 5.90 |
| Mutation | c.1067+1ins | cagAGTAAG | MAXENT: -12.23 | MM: -12.52 | WMM: -12.09 |
| Wild type | c.1067+1 | cagGTAAGC | MAXENT: 9.88 | MM: 11.18 | WMM: 11.85 |
| Mutation | c.2887+1C>T | gaaTTAAGT | MAXENT: 1.31 | MM: -0.41 | WMM: 0.20 |
| Wild type | c.2887+1C | gaaCTAAGT | MAXENT: 1.54 | MM: -0.18 | WMM: 0.43 |
| Mutation | c.746+3A>T | cctGTTAGT | MAXENT: -5.28 | MM: -6.27 | WMM: 1.66 |
| Wild type | c.746+3A | cctGTAAGT | MAXENT: 7.52 | MM: 6.68 | WMM: 6.61 |

| 3'splice site mutations | | | | | |
|---|---|---|---|---|---|
| | CDS location | Sequence | MAX-entropy | Markov Model | PWM |
| Mutation | c.1216-1G>A | gcattctgttgtgaccacaaGTG | MAXENT: -1.44 | MM: -0.36 | WMM: -2.01 |
| Wild type | c.1216-1G | gcattctgttgtgaccacagGTG | MAXENT: 7.31 | MM: 8.39 | WMM: 6.74 |
| Mutation | c.1504-8T>C | atcctctgttttcgctgtagGTT | MAXENT: 10.95 | MM: 10.57 | WMM: 12.40 |
| Wild type | c.1504-8T | atcctctgtttttgctgtagGTT | MAXENT: 10.78 | MM: 10.84 | WMM: 12.46 |
| Mutation | c.1216-6C>A | gcattctgttgtgaacacagGTG | MAXENT: 5.01 | MM: 6.48 | WMM: 4.10 |
| Wild type | c.1216-6C | gcattctgttgtgaccacagGTG | MAXENT: 7.31 | MM: 8.39 | WMM: 6.74 |
| Mutation | c.2561-1G>A | caatcatttgccatttctaaGAC | MAXENT: -0.13 | MM: -0.56 | WMM: -0.35 |
| Wild type | c.2561-1G>A | caatcatttgccatttctagGAC | MAXENT: 8.62 | MM: 8.19 | WMM: 8.40 |
| Mutation | c.585-1C>T | tctgcttttcattttattaaGCA | MAXENT: -0.59 | MM: -0.30 | WMM: 1.51 |
| Wild type | c.585-1C | tctgcttttcattttattagGCA | MAXENT: 8.16 | MM: 8.45 | WMM: 10.26 |
| Mutation | c.1603-1G>A | ctgtgtttttttattcccaaGAC | MAXENT: 1.30 | MM: 2.27 | WMM: 4.34 |
| Wild type | c.1603-1G | ctgtgtttttttattcccagGAC | MAXENT: 10.05 | MM: 11.02 | WMM: 13.09 |
| Mutation | c.655-1G>A | cttctttttaactttgacaaATA | MAXENT: -0.50 | MM: -0.77 | WMM: -0.26 |
| Wild type | c.655-1G | cttctttttaactttgacagATA | MAXENT: 8.25 | MM: 7.98 | WMM: 8.49 |

| Supplementary Table 4: TP53 mutations in BCC tumors | | | | |
|---|---|---|---|---|
| **Tumor-ID** | **Genomic position (NC_000017.10)** | **Exon / Intron** | **Mutation (NM_000546.5)** | **Amino acid (CCDS11118)** |
| 1 | g.7578401 | Exon 5 | c.529C>T | p.P177S |
| 4 | g.7577058 | Exon 8 | c.880G>T | p.E294* |
| 11 | g.7578212 | Exon 6 | c.637C>T | p.R213* |
| 13 | g.7578212 | Exon 6 | c.637C>T | p.R213* |
| | g.7577547_7577548 | Exon 7 | c.733_734GG>AA | p.G245N |
| 15 | g.7578400_7578401 | Exon 5 | c.529_530del18 | p.P177_C182 del PHHERC |
| | g.7577556 | Exon 7 | c.725G>T | p.C242F |
| 18 | g.7578395 | Exon 5 | c.535C>T | p.H179Y |
| | g.7577559 | Exon 7 | c.722C>T | p.S241F |
| 22 | g.7578395_7578396 | Exon 5 | c.534_535CC>TT | p.H178H; p.H179Y |
| 30 | g.7577124_7577125 | Exon 8 | c.813 814GG>AA | p.V272M |
| | g.7576855 | Exon 9 | c.991C>T | p.Q331* |
| 32 | g.7577532_7577533 | Exon 7 | c.748_749CC>TT | p.P250F |
| | g.7577082 | Exon 8 | c.856G>A | p.E286K |
| 33 | g.7578395 | Exon 5 | c.535C>T | p.H179Y |
| | g.7577046 | Exon 8 | c.892G>T | p.E298* |
| 46 | g.7578263 | Exon 6 | c.586C>T | p.R196* |
| 51 | g.7577106 | Exon 8 | c.832C>T | p.P278S |
| 54 | g.7577539 | Exon 7 | c.742C>T | p.R248T |
| | g.7577106 | Exon 8 | c.832C>T | p.P278S |
| 76 | g.7577559 | Exon 7 | c.722C>T | p.S241G |
| 123 | g.7578280 | Exon 6 | c.569C>T | p.P190L |
| 63 | g.7579313 | Exon 4 | c.374G>A | p.E67N |
| | g.7578472 | Exon 5 | c.458C>T | p.P153L |
| | g.7577079 | Exon 8 | c.859G>A | p.E287K |
| 64 | g.7578402 | Exon 5 | c.528C>G | p.C176T |
| | g.7577120 | Exon 8 | c.818G>A | p.R273H |
| 141 | g.7578275 | Exon 6 | c.574C>T | p.Q192* |
| 66 | g.7578399_7578400 | Exon 5 | c.530_531CC>TT | p.P177L |
| | g.7577531_7577532 | Exon 7 | c.749_750CC>TT | p.P250L |
| | g.7577559 | Exon 7 | c.722C>T | p.S241F |
| | g.7579313_7579335 | Intron 4 | c.375-23delAG | |
| 67 | g.7578517 | Exon 5 | c.413C>T | p.A138V |
| | g.7578263 | Exon 6 | c.586C>T | p.R196* |
| 7 | g.7578212 | Exon 6 | c.637C>T | p.R213* |
| 70 | g.7578212 | Exon 6 | c.637C>T | p.R213* |
| | g.7578263_7578264 | Exon 6 | c.585_586CC>TT | p.I195I; p.R196* |

| | | | | |
|---|---|---|---|---|
| | g.7577099 | Exon 8 | c.839G>A | p.R280K |
| 16 | g.7577509 | Exon 7 | c.772G>A | p.E258K |
| | g.7577099 | Exon 8 | c.839G>A | p.R280K |
| 19 | g.7578269 | Exon 6 | c.580C>T | p.L194F |
| | g.7578177_7578178 | Exon 6 | c.672G>A;c.672+1G>A | p.E224E |
| 34 | g.7578398 | Exon 5 | c.531_532CC>TG | p.H178D |
| 50 | g.7578253 | Exon 6 | c.596G>A | p.G199E |
| 81 | g.7578466 | Exon 5 | c.464C>T | p.T155S |
| | g.7578253 | Exon 6 | c.596C>T | p.G199V |
| | g.7578263 | Exon 6 | c.586C>T | p.R196* |
| | g.7577559 | Exon 7 | c.722C>T | p.S241F |
| 84 | g.7578471 | Exon 5 | c.459delC | p.G154fs*16 |
| | g.7577102 | Exon 8 | c.836G>A | p.G279E |
| | g.7577106 | Exon 8 | c.832C>T | p.P278S |
| | g.7576897_7576898 | Exon 9 | c.948_949GG>AA | p. Q317* |
| 85 | g.7578400 | Exon 5 | c.530C>T | p.P177L |
| | g.7578475_7578476 | Exon 5 | c.454_455CC>TT | p.P152L |
| | g.7578175 | Intron 6 | c.672+2A>T | |
| 90 | g.7577099 | Exon 8 | c.839G>T | p.R280I |
| 91 | g.7577574 | Exon 7 | c.707A>G | p.Y236C |
| 94 | g.7578275 | Exon 6 | c.574C>T | p.Q192* |
| | g.7577547_7577548 | Exon 7 | c.733_734GG>TT | p.G245F |
| 95 | g.7577539_7577540 | Exon 7 | c.741_742CC>TT | p.N247N; R248T |
| 98 | g.7578524 | Exon 5 | c.406C>T | p.Q136* |
| 102 | g.7578212 | Exon 6 | c.637C>T | p.R213* |
| | g.7577559 | Exon 7 | c.722C>T | p.S241G |
| 104 | g.7577547 | Exon 7 | c.734G>A | p.G245D |
| 105 | g.7578400_7578401 | Exon 5 | c.530_531CC>TT | p.P177L |
| | g.7577082 | Exon 8 | c.856G>A | p.E286K |
| 106 | g.7578400 | Exon 5 | c.530C>T | p.P177L |
| | g.7577137_7577138 | Exon 8 | c.800_801GG>AA | p.R267Q |
| 111 | g.7578212 | Exon 6 | c.637C>T | p.R213* |
| 119 | g.7577509_7577510 | Exon 7 | c.771_772GG>AA | p.L257L; p.E258K |
| 132 | g.7578395 | Exon 5 | c.535C>T | p.H179Y |
| | g.7577159 | Intron 7 | c.783-4 G>A | |
| 134 | g.7577538 | Exon 7 | c.743G>A | p.R248P |
| 139 | g.7578400 | Exon 5 | c.530C>T | p.P177L |
| | g.7577547_7577548 | Exon 7 | c.733_734GG>TT | p.G245F |
| 146 | g.7577559 | Exon 7 | c.722C>T | p.S241G |
| | g.7577094 | Exon 8 | c.844C>T | p.R282T |

| | | | | |
|---|---|---|---|---|
| 147 | g.7577537_7577538 | Exon 7 | c.743_744GG>AA | p.R248Q |
| 148 | g.7578181 | Exon 6 | c.668C>T | p.P223L |
| | g.7577142_7577143 | Exon 8 | c.795_796GG>AA | p.L265L; p.G266R |
| 150 | g.7578398 | Exon 5 | c.532C>T | p.H178Y |
| 151 | g.7578513 | Exon 5 | c.417G>C | p.K139N |
| | g.7578549_7578550 | Exon 5 | c.380_381CC>TT | p.S127S; p.P128S |
| 153 | g.7578524 | Exon 5 | c.406C>T | p.Q136* |
| 160 | g.7578400 | Exon 5 | c.530C>T | p.P177L |
| | g.7578212 | Exon 6 | c.637C>T | p.R213* |
| 165 | g.7578212 | Exon 6 | c.637C>T | p.R213* |
| | g.7577102 | Exon 8 | c.836G>A | p.G279E |
| 168 | g.7578263 | Exon 6 | c.586C>T | p.R196* |
| | g.7577082 | Exon 8 | c.856G>T | p.E286* |
| 170 | g.7578395 | Exon 5 | c.535C>T | p.H179Y |
| 174 | g.7577575 | Exon 7 | c.706T>C | p.Y236H |
| 175 | g.7578400 | Exon 5 | c.530C>T | p.P177L |
| | g.7578550 | Exon 5 | c.380C>T | p.S127F |
| 185 | g.7578461 | Exon 5 | c.469G>T | p.V157F |
| | g.7577498 | Intron 7 | c.782+1G>A | |
| 186 | g.7576899 | Exon 9 | c.947C>T | p.P316L |
| 187 | g.7577547 | Exon 7 | c.734G>A | p.G245D |
| 189 | g.7578400_7578401 | Exon 5 | c.530_531CC>TT | p.P177L |
| 190 | g.7578550 | Exon 5 | c.380C>T | p.S127F |

| Supplementary Table 5: Univariate analysis for the effect of mutations in PTCH1, TP53, TERT promoter and DPH3 promoter on patient characteristics | | | | | | | | | | | | | | | | |
|---|---|---|---|---|---|---|---|---|---|---|---|---|---|---|---|---|
| | **PTCH1** | | | | **TP53** | | | | **TERT promoter** | | | | **DPH3 promoter** | | | |
| | **Wt** | **Mut** | **OR (95% CI)** | **P** | **Wt** | **Mut** | **OR (95% CI)** | **P** | **Wt** | **Mut** | **OR** | **P** | **Wt** | **Mut** | **OR** | **P** |
| **Age** | | | | | | | | | | | | | | | | |
| <70 | 31 | 58 | Reference | 0.07 | 57 | 32 | Reference | 0.60 | 42 | 47 | Reference | 0.10 | 61 | 28 | Reference | 0.07 |
| >70 | 49 | 53 | 0.58 (0.32-1.04) | | 69 | 33 | 0.85 (0.47-1.55) | | 36 | 66 | 1.64 (0.92-2.93) | | 56 | 45 | 1.75 (0.97-3.17) | |
| **Gender** | | | | | | | | | | | | | | | | |
| Men | 50 | 65 | Reference | 0.58 | 71 | 44 | Reference | 0.13 | 49 | 66 | Reference | 0.54 | 69 | 45 | Reference | 0.71 |
| Women | 30 | 46 | 1.18 (0.65-2.13) | | 55 | 21 | 0.61 (0.33-1.15) | | 29 | 47 | 1.20 (0.67-2.18) | | 48 | 28 | 0.89 (0.49-1.63) | |
| **Fototype** | | | | | | | | | | | | | | | | |
| 3 to 5 | 48 | 54 | Reference | | 74 | 28 | Reference | | 48 | 54 | Reference | | 66 | 35 | Reference | |
| 1 or 2 | 27 | 47 | 1.55 (0.84-2.86) | 0.16 | 41 | 33 | **2.13 (1.13-4.00)** | **0.02** | 25 | 49 | 1.74 (0.94-3.24) | 0.08 | 42 | 32 | 1.50 (0.81-2.79) | 0.55 |
| *Not stated* | 5 | 10 | | | 11 | 4 | | | 5 | 10 | | | 8 | 7 | | |
| **Nevus count** | | | | | | | | | | | | | | | | |
| < 50 | 63 | 80 | Reference | 0.11 | 98 | 45 | Reference | 0.04 | 55 | 88 | Reference | 0.33 | 88 | 54 | Reference | 0.49 |
| > 50 | 5 | 15 | 2.36 (0.82-6.85) | | 9 | 11 | **2.66 (1.03-6.87)** | | 10 | 10 | 0.63 (0.24-1.60) | | 14 | 6 | 0.70 (0.25-1.93) | |
| *Not stated* | 12 | 16 | | | 19 | 9 | | | 13 | 15 | | | 15 | 13 | | |
| **Presence of solar lentigos** | | | | | | | | | | | | | | | | |
| No | 6 | 13 | Reference | 0.52 | 11 | 8 | Reference | 0.46 | 8 | 11 | Reference | 0.79 | 11 | 8 | Reference | 0.51 |
| Yes, scarce | 28 | 41 | 0.67 (0.23-1.99) | | 49 | 20 | 0.56 (0.20-1.60) | | 30 | 39 | 0.95 (0.34-2.64) | | 40 | 29 | 1.00 (0.36-2.79) | |
| Yes, abundant | 36 | 43 | 0.55 (0.19-1.60) | | 50 | 29 | 0.80 (0.29-2.21) | | 30 | 49 | 1.19 (0.43-3.29) | | 52 | 26 | 0.69 (0.25-1.92) | |
| *Not stated* | 10 | 14 | | | 16 | 8 | | | 10 | 14 | | | 14 | 10 | | |
| **History of cutaneous neoplasms** | | | | | | | | | | | | | | | | |
| No | 34 | 33 | Reference | 0.09 | 47 | 20 | Reference | 0.40 | 36 | 31 | Reference | 0.04 | 49 | 18 | Reference | 0.03 |

| | | | | | | | | | | | | | | | | |
|---|---|---|---|---|---|---|---|---|---|---|---|---|---|---|---|---|
| Yes | 27 | 47 | 1.79 (0.92-3.52) | | 47 | 27 | 1.35 (0.67-2.73) | | 27 | 47 | **2.02 (1.03-3.97)** | | 40 | 33 | **2.25 (1.10-4.57)** | |
| *Not stated* | 19 | 31 | | | 32 | 18 | | | 15 | 35 | | | 28 | 22 | | |
| **Sun-exposure** | | | | | | | | | | | | | | | | |
| None | 42 | 54 | Reference | 0.45 | 62 | 34 | Reference | 0.24 | 37 | 59 | Reference | 0.01 | 62 | 33 | Reference | 0.09 |
| <20y | 16 | 14 | 0.68 (0.30-1.55) | | 23 | 7 | 0.56 (0.22-1.43) | | 20 | 10 | **0.31 (0.13-0.74)** | | 22 | 8 | 0.68 (0.27-1.70) | |
| >20y | 14 | 23 | 1.28 (0.59-2.78) | | 21 | 16 | 1.39 (0.64-3.01) | | 11 | 26 | 1.48 (0.66-3.35) | | 18 | 19 | 1.98 (0.92-4.29) | |
| *Not stated* | 8 | 20 | | | 20 | 8 | | | 10 | 18 | | | 15 | 13 | | |
| **Immuno-supression** | | | | | | | | | | | | | | | | |
| No | 66 | 94 | Reference | 0.15 | 105 | 55 | Reference | 0.95 | 64 | 96 | Reference | 0.36 | 98 | 61 | Reference | 0.34 |
| Yes | 6 | 3 | 0.35 (0.09-1.45) | | 6 | 3 | 0.96 (0.23-3.97) | | 5 | 4 | 0.53 (0.14-2.06) | | 7 | 2 | 0.46 (0.09-2.28) | |
| *Not stated* | 8 | 14 | | | 15 | 7 | | | 9 | 13 | | | 12 | 10 | | |
| **Risk type (based on histology)** | | | | | | | | | | | | | | | | |
| Low risk | 43 | 73 | Reference | 0.98 | 73 | 43 | Reference | 0.54 | 43 | 73 | Reference | 0.23 | 72 | 43 | Reference | 0.42 |
| High risk | 14 | 24 | 1.01 (0.47-2.16) | | 26 | 12 | 0.78 (0.36-1.71) | | 10 | 28 | 1.65 (0.73-3.72) | | 21 | 17 | 1.36 (0.65-2.85) | |
| *Not stated* | 23 | 14 | | | 27 | 10 | | | 25 | 12 | | | 24 | 13 | | |

| Supplementary Table 6: PCR Conditions and Primer Sequences | | | | |
|---|---|---|---|---|
| **Name** | **Forward primer** | **Reverse primer** | **Tm** | **Product size (bp)** |
| TERT promoter: -27 to -286 | 5'CCCACGTGCGCAGCAGGAC 3' | 5'CTCCCAGTGGATTCGCGGGC 3' | 60°C | 260 |
| DPH3-OXNAD1 promoter | 5'CGAAGGGGTAACGCCCCAG 3' | 5'GGTCCCAGACGTGACGTAGC 3' | 56°C | 314 |
| TP53: exon 5/6 | 5'CTGCCGTCTTCCAGTTG 3' | 5'CTGACAACCACCCTTAACC 3' | 50°C | 504 |
| TP53: exon 7 | 5'AAGGCGCACTGGCCTCATCTT 3' | 5'AGTGGGAGCAGTAAGGAGATT 3' | 56°C | 307 |
| TP53: exon 8/9 | 5'CTTACTGCCTCTTGCTTCTC 3' | 5'CCACTTGATAAGAGGTCCC 3' | 50°C | 373 |
| PTCH1: exon 4 | 5'TCCAGGGCAACTTCATTTACTA 3' | 5'CCAGTCTCATGACTAGAGTTCA 3' | 54°C | 410 |
| PTCH1: exon 5/6 | 5'CATTCTGCTGAAATCCCCTCT 3' | 5'TACTTGGCAAAAGCTCTGCT 3' | 54°C | 571 |
| PTCH1: exon 7 | 5'TGCTCTCCACCCTTCTGAGAG 3' | 5'CTCCTACAAGGTGGATGCAGTG 3' | 57°C | 335 |
| PTCH1: exon 8 | 5'GCTAGCGAGGATAACGGTTTAAG 3' | 5'GCATGTGACCTGCCTACTAATTC 3' | 55°C | 296 |
| PTCH1: exon 9 | 5'TGCATAACCAGCGAGTCTGC 3' | 5'GAATTGCAGCCAGTGAGTTGG 3' | 57°C | 295 |
| PTCH1: exon 10 | 5'CCTGGATGCACATCGATGTT 3' | 5'TGTCGAGGCTTGTGGAAGT 3' | 56°C | 308 |
| PTCH1: exon 11 | 5'GTAGCATCCTAGTGGAAAAGGC 3' | 5'TGATGGGTGGAGGGAAACATTA 3' | 54°C | 375 |
| PTCH1: exon 12 | 5'CTTAGGAACAGAGGAAGCTGTGA 3' | 5'CTGCTTCAGGAGCTGTTAGGT 3' | 54°C | 265 |
| PTCH1: exon 13 | 5'CAGAGCCTCAAACACAGGCA 3' | 5'TCACACAGTCTGTGCTCCAAG 3' | 56°C | 329 |
| PTCH1: exon 14/15 | 5'CTCCCATGGAAGATGACCTCA 3' | 5'AGGCCCAGAAACAGTCAAGAC 3' | 57°C | 1582 |
| PTCH1: exon 16 | 5'CTCCACCTGTTACAACATGCTC 3' | 5'CTATGGGAGAACAACCCCTACA 3' | 54°C | 726 |
| PTCH1: exon 17 | 5'TGCCTTAGGTCTCCAGAGAGCA 3' | 5'GCTAGGACCAGGGTCCTTCT 3' | 56°C | 318 |
| PTCH1: exon 18 | 5'GTAGCGTCAACGGATGAAGG 3' | 5'GTAATGCTGTGCGAAGCTCAG 3' | 54°C | 375 |
| PTCH1: exon 19 | 5'ATGCAAGCTATACCCTCCTCC 3' | 5'GAGCAGTCAGTAAGCTTTCATGC 3' | 56°C | 497 |
| PTCH1: exon 20 | 5'ACTTGGAGACAAACAGAGCCA 3' | 5'ATCTGAACCGAGGACACCTTA 3' | 54°C | 310 |
| PTCH1: exon 21 | 5'ACCCGGCCCAATCACAATGATT 3' | 5'AGATGTGAGCAGTTCTGAGAGC 3' | 57°C | 334 |
| PTCH1: exon 22 | 5'ACTGAAGAACCACCAGCAAGTG 3' | 5'TGTTCATTTCTGGCGTTGCCAT 3' | 57°C | 314 |
| PTCH1: exon 23 | 5'TGTGTGATGTGCTGCTCAGCA 3' | 5'CTTTGAGTGTGGCCAGCAGGTA 3' | 59.5°C | 412 |
| PTCH1: exon 24 | 5'GACAAAGCTTGGACACATCAGC 3' | 5'CTGAGTCTTTGGTGAAACCCA 3' | 56.5°C | 812 |

#### Supplementary Figure 1

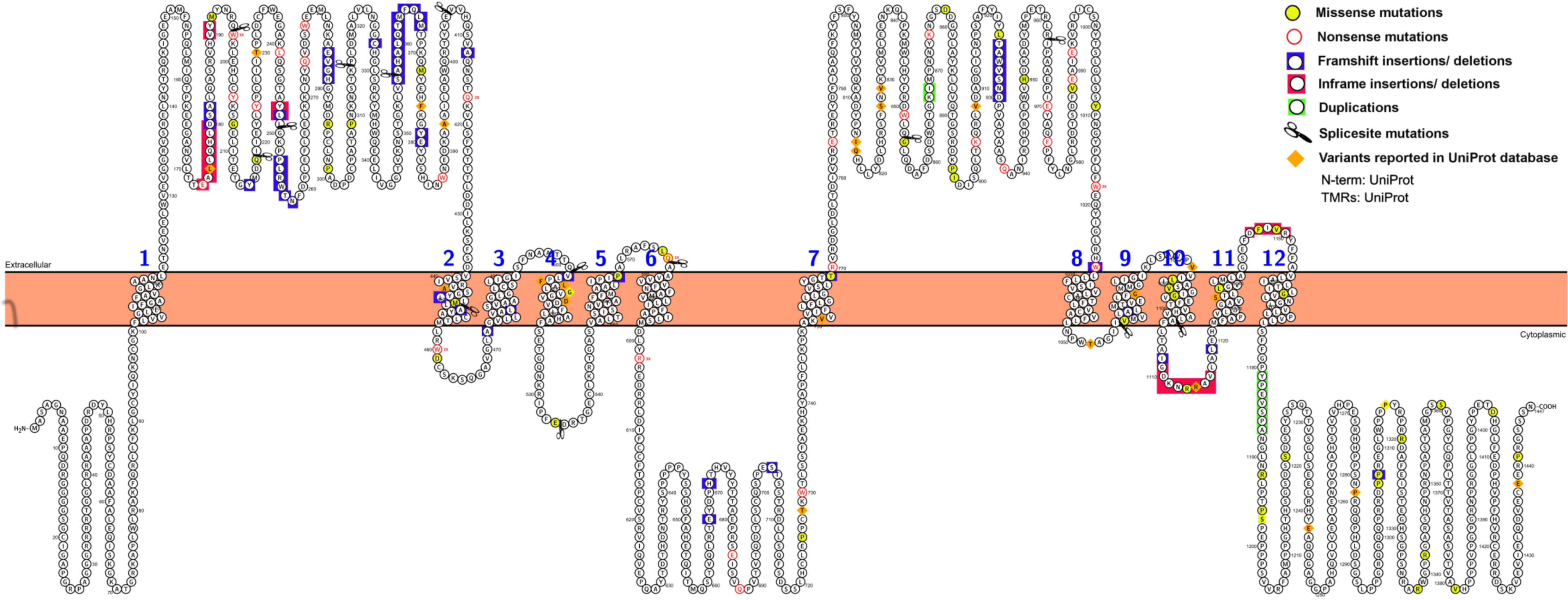

#### Supplementary Figure 2

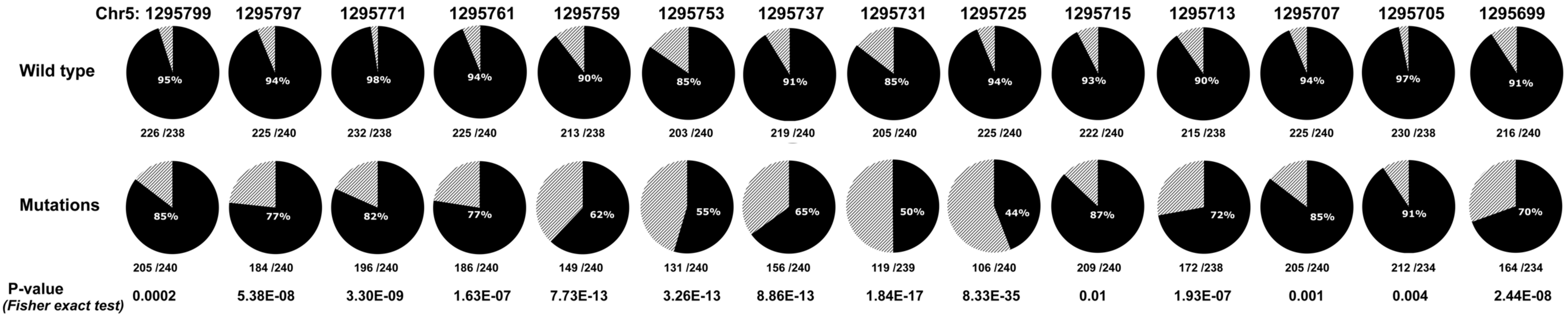

**Supplementary Figure 3**

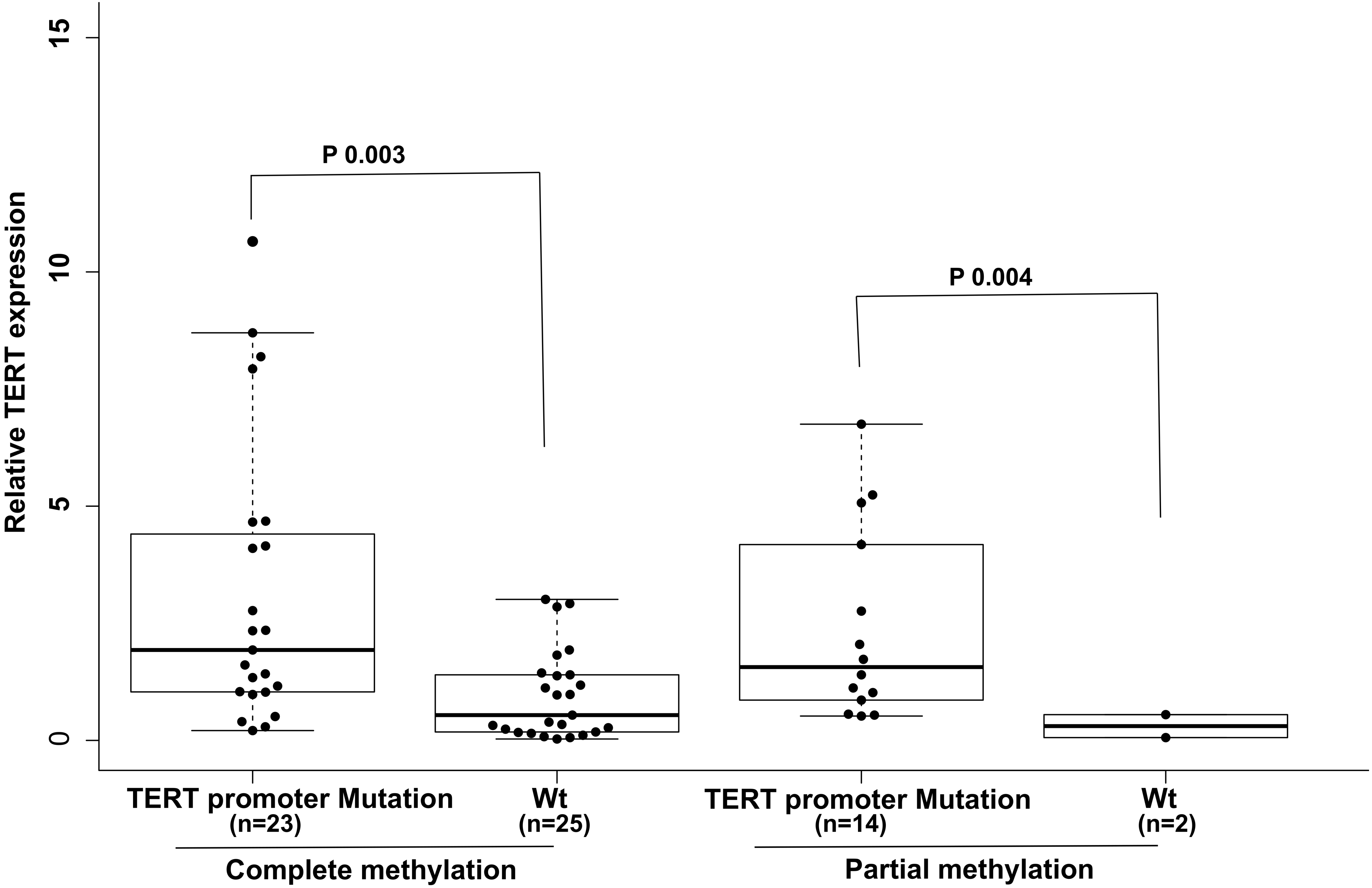

### Supplementary Figure 4

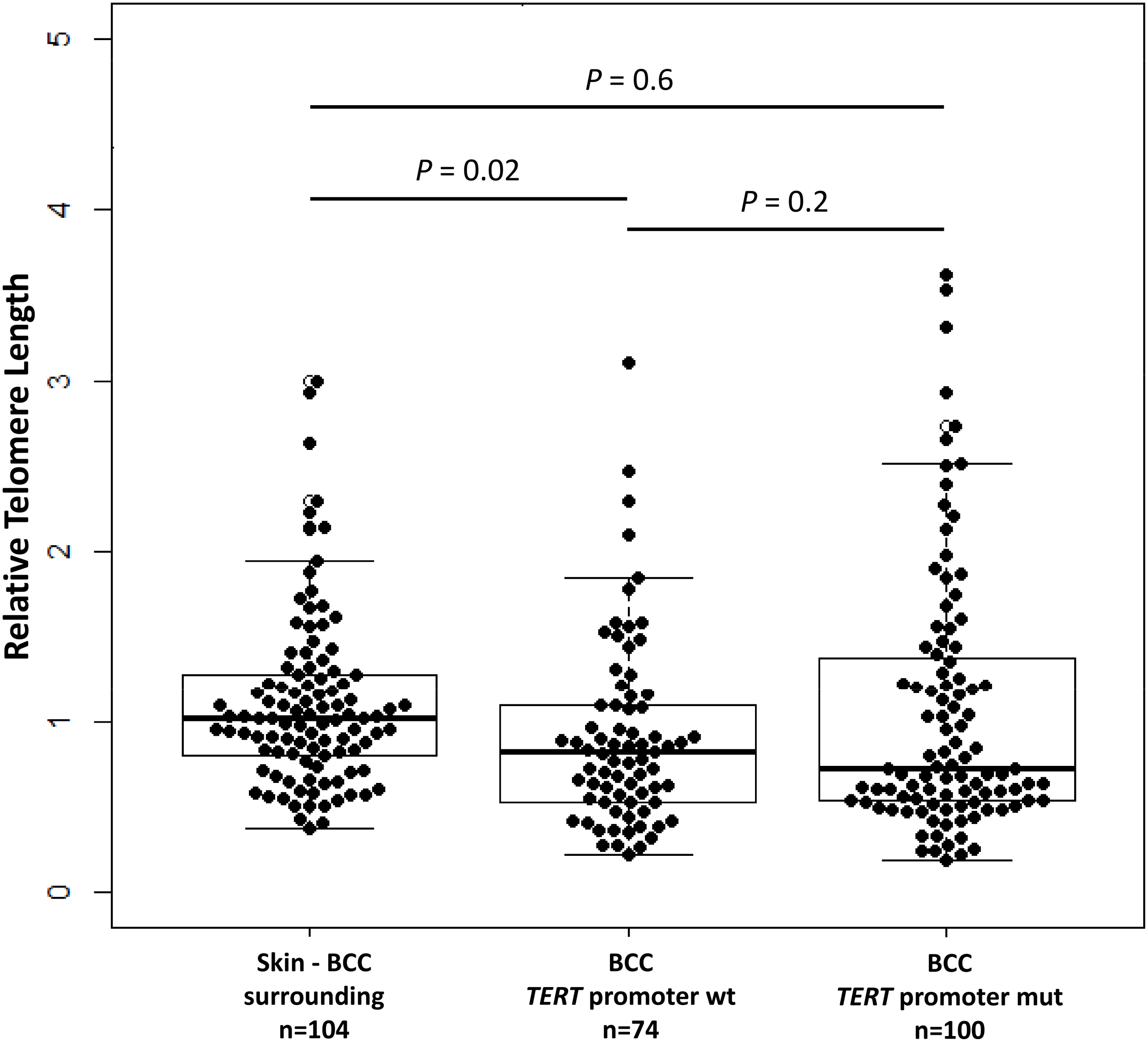
